# Brown adipocyte fatty acid synthase (FASN) deficiency protects mice from alcohol-induced elevations in plasma triglyceride and hepatic steatosis

**DOI:** 10.64898/2026.08.22.746452

**Authors:** Preethi Parupalli, Piumi Wickramasinghe, Laura Hua, Lin Jia

## Abstract

Excessive alcohol intake is frequently associated with hypertriglyceridemia, a condition that increases the risk of severe complications including acute pancreatitis and cardiovascular disease. The very low-density lipoprotein (VLDL) receptor (VLDLR) promotes uptake of apoE-containing VLDL particles by peripheral tissues and plays an important role in maintaining plasma triglyceride (TG) homeostasis. Brown adipose tissue (BAT) is a major metabolic organ that contributes to circulating lipid clearance during thermogenic activation. It was reported that cold-induced thermogenesis upregulates VLDLR expression in BAT and reduces plasma TG via VLDL uptake. However, whether BAT VLDLR-mediated VLDL uptake regulates alcohol-induced hypertriglyceridemia remains unknown. Here, we generated BAT-specific fatty acid synthase (FASN) knockout mice (FASN^BKO^) and subjected them to binge and acute-on-chronic alcohol feeding paradigms. We found that BAT FASN deficiency enhanced thermogenic function and promoted VLDL uptake, resulting in attenuation of alcohol-induced elevations in plasma TG. Consistent with these findings, pharmacological inhibition of FASN by TVB3664 treatment in differentiated brown adipocytes (bADs) increased thermogenic gene expression and VLDL uptake under both control and alcohol-exposed conditions. In addition, FASN^BKO^ mice were protected from alcohol-induced hepatic steatosis, which was accompanied by increased hepatic AMP-activated-protein kinase (AMPK) activation and enhanced β-oxidation. Furthermore, FASN^BKO^ mice exhibited upregulated FGF21 mRNA expression in the BAT and elevated circulating FGF21 levels. Similarly, TVB3664-treated differentiated bADs showed higher FGF21 expression and increased FGF21 content in culture medium. Taken together, these findings identify the important role of brown adipocyte FASN in regulating thermogenic function and TG homeostasis during alcohol exposure and suggest that enhancing thermogenic lipid utilization in BAT may represent a potential therapeutic strategy for mitigating alcohol-associated increases in plasma TG and hepatic fat accumulation.

**SUMMARY:** Brown adipose tissue (BAT)-specific fatty acid synthase (FASN) deletion enhances triglyceride-rich very low-density lipoprotein clearance during alcohol exposure, leading to reduced plasma triglyceride (TG) in mice. BAT-specific FASN deficiency also protects mice from alcohol-induced hepatic steatosis probably through BAT-liver metabolic signaling.

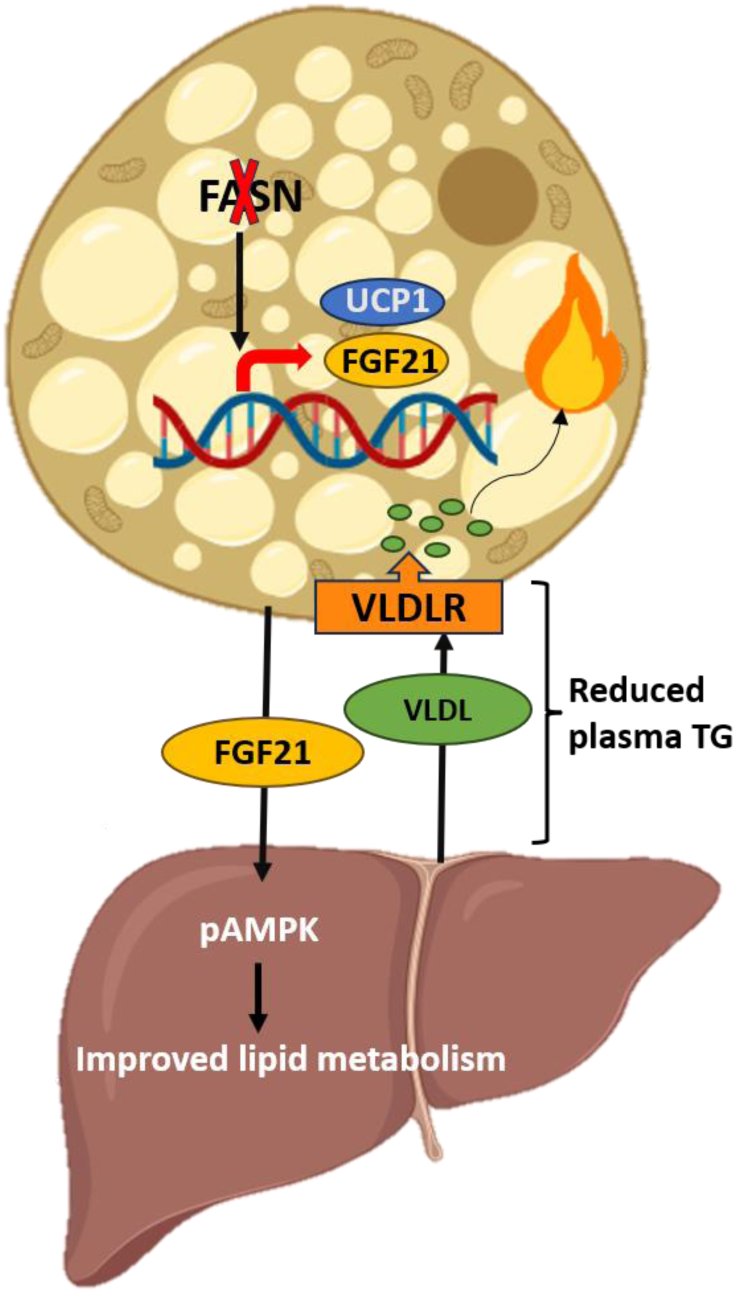

## Introduction

Excessive alcohol consumption is a major global health concern and contributes substantially to morbidity and mortality worldwide. Alcohol use was responsible for approximately 2.6 million deaths globally in 2019, reflecting its widespread impact on multiple organ systems ^1^. In addition to the development of alcohol-associated liver disease (ALD), one of the most recognized consequences of excessive alcohol exposure, alcohol overconsumption disrupts systemic lipid homeostasis and is associated with increased circulating triglyceride (TG) levels in humans and experimental models ^2–5^. Alcohol-induced increases in plasma TG is clinically important because severe elevation of circulating TG levels is a major risk factor for acute pancreatitis ^6–8^. Alcohol itself is also an important secondary cause of both hypertriglyceridemia and pancreatitis ^7^. Although several TG-lowering strategies are available, there are currently no approved therapies specifically targeting alcohol-induced hypertriglyceridemia. Therefore, identifying the mechanisms responsible for alcohol-induced disruption of TG homeostasis is essential for developing targeted therapeutic strategies.

Plasma TG concentrations are determined by the balance between hepatic production of triglyceride-rich lipoproteins (TRLs) and their clearance by peripheral tissues. Following synthesis in hepatocytes, TG are packaged into very low-density lipoproteins (VLDL) and secreted into the circulation, where they serve as the primary vehicle for lipid transport to metabolically active tissues ^9^. Circulating TG-rich lipoproteins, including VLDL and intestine-derived chylomicrons, are classically cleared through lipoprotein lipase (LPL)-mediated hydrolysis on capillary endothelial surfaces in adipose tissue, skeletal muscle, and BAT ^10–12^. LPL hydrolyzes TG within these particles, releasing free fatty acids (FFA) for tissue uptake, storage, or oxidation ^13^. Therefore, disruption of either hepatic VLDL secretion or peripheral TG clearance can result in elevated plasma TG levels ^14,15^. This concept is supported by the findings that chronic alcohol exposure promotes hepatic VLDL production, which is accompanied by increased plasma TG and that LPL deficient mice display markedly impaired TG-rich lipoprotein catabolism and severe hypertriglyceridemia ^16,17^

Zemánková et al. reported that acute alcohol exposure in humans downregulates LPL activity ^18^. In addition, Lecomte et al. found that humans drinking excessive alcohol exhibit increased concentration of circulating apoC-III, a potent inhibitor of LPL ^19^. However, LPL suppression alone does not fully explain alcohol-induced hypertriglyceridemia ^3,20–22^. Of note, recent findings indicate the important role of VLDL receptor (VLDLR)-mediated uptake of VLDL particles in circulating TG clearance in cold-exposed mice. Specifically, Shin et al. reported that cold-induced thermogenesis is associated with upregulated expression of VLDLR in BAT and brown adipocytes (bADs), facilitating the uptake of circulating VLDL-derived lipids ^23^. This enhanced lipid uptake provides essential fuel for adaptive thermogenesis and contributes to the reduction of plasma TG levels. However, it remains unclear whether enhancing thermogenic program in BAT can regulate TG homeostasis during alcohol exposure.

Fatty acid synthase (FASN), a key enzyme involved in de novo lipogenesis, catalyzes the production of saturated long-chain fatty acids from acetyl-CoA and malonyl-CoA. Several studies using white adipose tissue (WAT) and cultured white adipocytes showed that adipocyte-specific FASN deletion or inhibition leads to uncoupling protein 1 (UCP1) upregulation and increased thermogenic function ^24–28^. Lodhi et al. reported that constitutive adipocyte-specific FASN ablated mice exhibit enhanced thermogenic programming, including UCP1 induction in inguinal WAT (iWAT), increased energy expenditure, and protection against diet-induced obesity ^25^. Similarly, using inducible adipocyte-specific FASN knockout mice maintained on a standard chow diet, Rowland et al. and Guilherme et al. demonstrated that loss of adipocyte FASN promotes UCP1 induction and beige adipocyte formation in iWAT ^26,28^. Regarding the effect of FASN deficiency on BAT thermogenic function, two studies found that BAT-specific FASN deletion could not elevate the expression of UCP1 ^26,28^. In contrast, a recent finding by Korobkina et al. showed that mice lacking FASN specifically in brown adipocytes display increased UCP1 protein levels in mouse BAT ^27^. Nevertheless, whether BAT-specific FASN deficiency alters thermogenic program in alcohol-fed mice remains unknown.

Recent studies have demonstrated that activation of thermogenic adipose tissue can protect against alcohol-induced metabolic dysfunction and liver injury. Shen et al. found that, in chronic ethanol-fed C57BL/6J mice, cold-induced BAT activation enhances UCP1-dependent mitochondrial thermogenesis, thereby attenuating ethanol-induced hepatic steatosis and liver injury. In line with these findings, Fan et al. showed that pharmacological activation of BAT thermogenic activity through bile acid signaling protects mice from alcohol-induced liver damage ^29^. In contrast, inhibition of thermogenic function in adipose tissues abolishes these protective effects and markedly exacerbates ALD ^29–31^. However, whether enhanced thermogenic activity promotes peripheral TG clearance and prevents alcohol-induced hypertriglyceridemia remains unknown. In the current study, using a BAT-specific FASN knockout model and pharmacological inhibition of FASN in bADs, we found that loss of FASN enhances thermogenic function, promotes VLDLR-mediated VLDL clearance, and protects against alcohol-induced increases in plasma TG during binge and acute-on-chronic ethanol exposure.

## Results

### BAT FASN deletion enhances BAT thermogenesis in control and acute alcohol-fed mice

Mice lacking FASN in BAT (FASN^BKO^) were generated and dramatically reduced FASN expression was observed at both the transcript and protein levels in chow-fed mice (**Supplementary Figure 1A–B**). It was reported that brown adipocyte FASN deficiency was associated with elevated UCP1 expression in the BAT ^27^. To investigate whether brown adipocyte FASN regulates thermogenic program in alcohol-fed mice, male FANS^fl/fl^ and FASN^BKO^ mice were subjected to binge drinking. Compared to FASN^fl/fl^ mice, FASN^BKO^ mice exhibited visibly darker interscapular BAT under both control and acute alcohol-treated conditions (**Figure 1A**), which was accompanied by significantly increased UCP1 levels examined by qPCR and Western blotting (**Figure 1B-C**). Similar findings were observed by immunostaining BAT with UCP1 (**Figure 1E**). Consistent with these observations, mRNA levels of key thermogenic genes, including Pgc1α, Pparγ, and Prdm16 were significantly elevated in the BAT of FASN^BKO^ relative to FASN^fl/fl^ mice under both feeding conditions (**Figure 1D**). In parallel, protein levels of mitochondrial markers (TOM20 and CPT1a) were significantly increased, indicating enhanced mitochondrial abundance and respiratory capacity (**Figure 1F**). Interestingly, mRNA expression of Cidea was significantly reduced in both control-and alcohol-fed FASN^BKO^ mice (**Figure 1G**). Binge drinking caused an increase in mRNA expression (**Figure 1B**), but not protein levels of UCP1 (**Figure 1C**) in FASN^fl/fl^ mice. Compared to control-fed FASN^fl/fl^ mice, acute alcohol-treated FASN^fl/fl^ mice exhibited more lipid droplets and significantly higher TG content in the BAT (**Figure 1H and 1I**). Of note, lipid accumulation was markedly reduced in the BAT of FASN^BKO^ relative to FASN^fl/fl^ mice following acute alcohol exposure (**Figure 1H and 1I**). Previous studies have shown that BAT FASN knockdown leads to increases in protein levels of p62 ^28^. In line with this, higher p62 expression was observed in the BAT of both control-and alcohol-fed FASN^BKO^ mice (**Supplementary Figure 2A-B**). Chen et al. reported that chronic alcohol feeding suppresses FASN expression in WAT ^32^. Consistently, we found slightly reduced FASN expression in both gonadal WAT (gWAT) and iWAT of acute alcohol-fed FASN^fl/fl^ mice compared to FASN^fl/fl^ mice fed control diet (**Supplementary Figure 1C**). Interestingly, the FASN expression in WAT of FASN^BKO^ was not regulated by acute alcohol treatment. In addition, FASN expression in WAT was comparable between FASN^fl/fl^ and FASN^BKO^ mice, indicating the specific gene deletion in BAT mediated by UCP1-Cre (**Supplementary Figure 1C**). Together, these findings suggest that loss of FASN in brown adipocytes promotes thermogenic program and limits lipid deposition in BAT.

**Figure 1.**
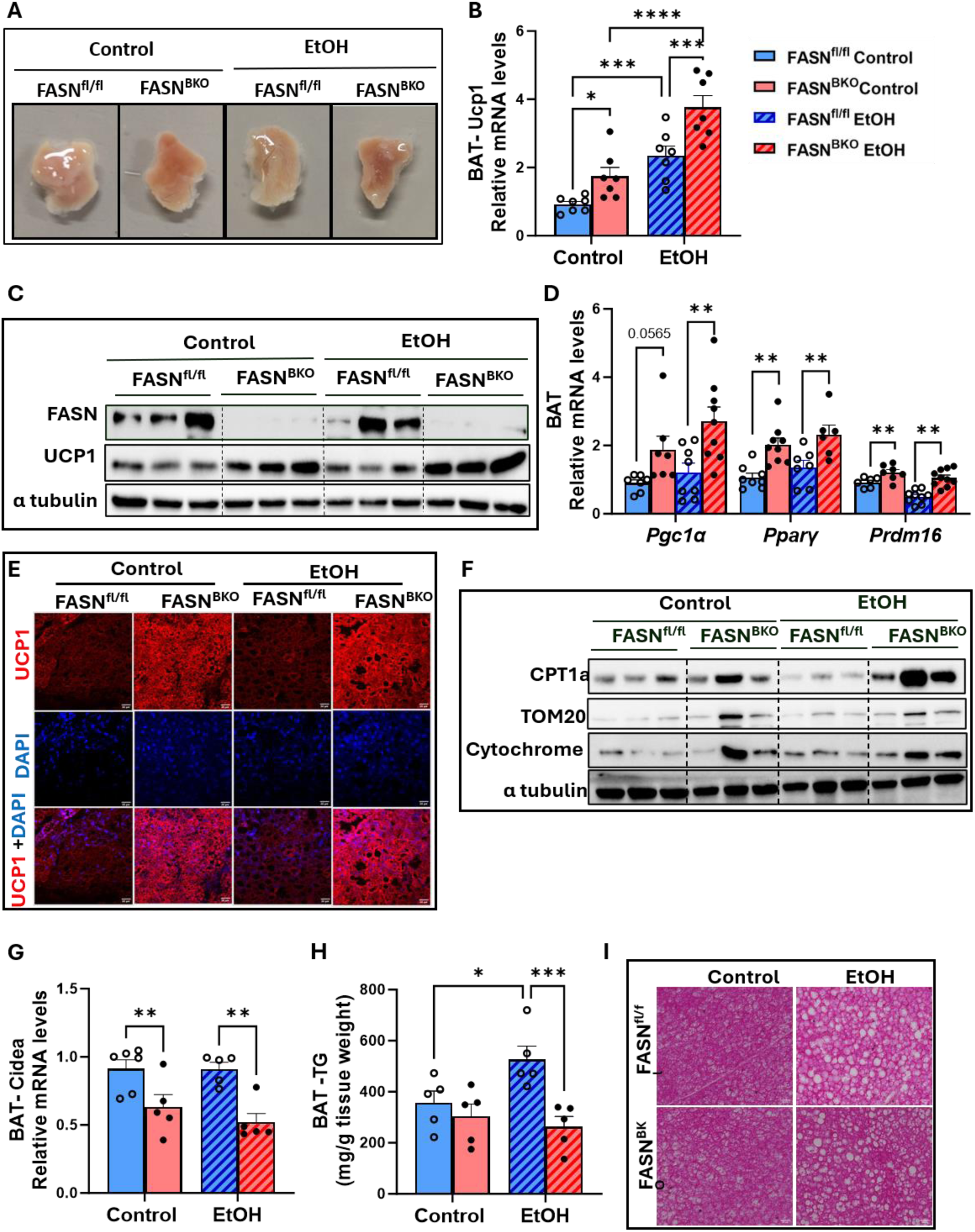
BAT-specific FASN deletion induces thermogenesis. Male FASN^fl/fl^ and FASN^BKO^ mice were subjected to a single alcohol binge (EtOH, 5 g/kg BW) or isocaloric maltose dextrin (control) via oral gavage. (A) Representative images of BAT tissues. (B) qPCR analysis of mRNA levels of UCP1 in BAT (n=7). (C) Western blotting of FASN, UCP1 and α tubulin in BAT. (D) qPCR analysis of PGC1α, PPARγ and Prdm16 in BAT (n=6-8). (E) Immunofluorescent staining of BAT UCP1. (F) Western blotting of CPT1a, TOM20, Cytochrome c, and α tubulin in BAT. (G) qPCR analysis of Cidea in BAT (n= 5). (H) BAT TG content (n=5). (I) Representative images of H&E staining of BAT tissues. Data are expressed as the mean ± SEM. *P <.05, **P <.01, ***P <.001,**** P<.0001.

### Pharmacological inhibition of FASN in cultured brown adipocytes upregulates thermogenic gene expression

Next, we performed in vitro assays using differentiated bADs, which were treated with either vehicle or TVB3664 (100nM, a FASN inhibitor) in the presence or absence of alcohol (100mM). Similar to the findings reported by Wan et al.^33^, we observed a reduction in both the number and size of lipid droplets in TVB3664-treated mature bADs following both control and EtOH exposure (**Figure 2A**). Consistent with our in vivo findings, mRNA expression of thermogenic genes such as Ucp1, Pgc1α, Cpt1a, Acox1 and Prdm16 were significantly elevated in TVB3664-treated bADs (**Figure 2B**). In addition, **Figure 2C** showed elevated UCP1 immunostaining in TVB3664-treated bADs under both control and EtOH conditions. Similar findings were observed after a lower dose of EtOH exposure (35mM) (**Supplementary Figure 3A**). Rowland et al. reported that TVB3664-mediated FASN inhibition causes elevated p62 expression in cultured primary white adipocytes^28^. In agreement, increased p62 expression was observed in TVB3664-treated bADs following both control and alcohol exposure (**Supplementary Figure 3B**). Thus, our in vitro data indicates that FASN inhibition in cultured bADs leads to upregulated thermogenesis irrespective of control or alcohol exposure.

**Figure 2.**
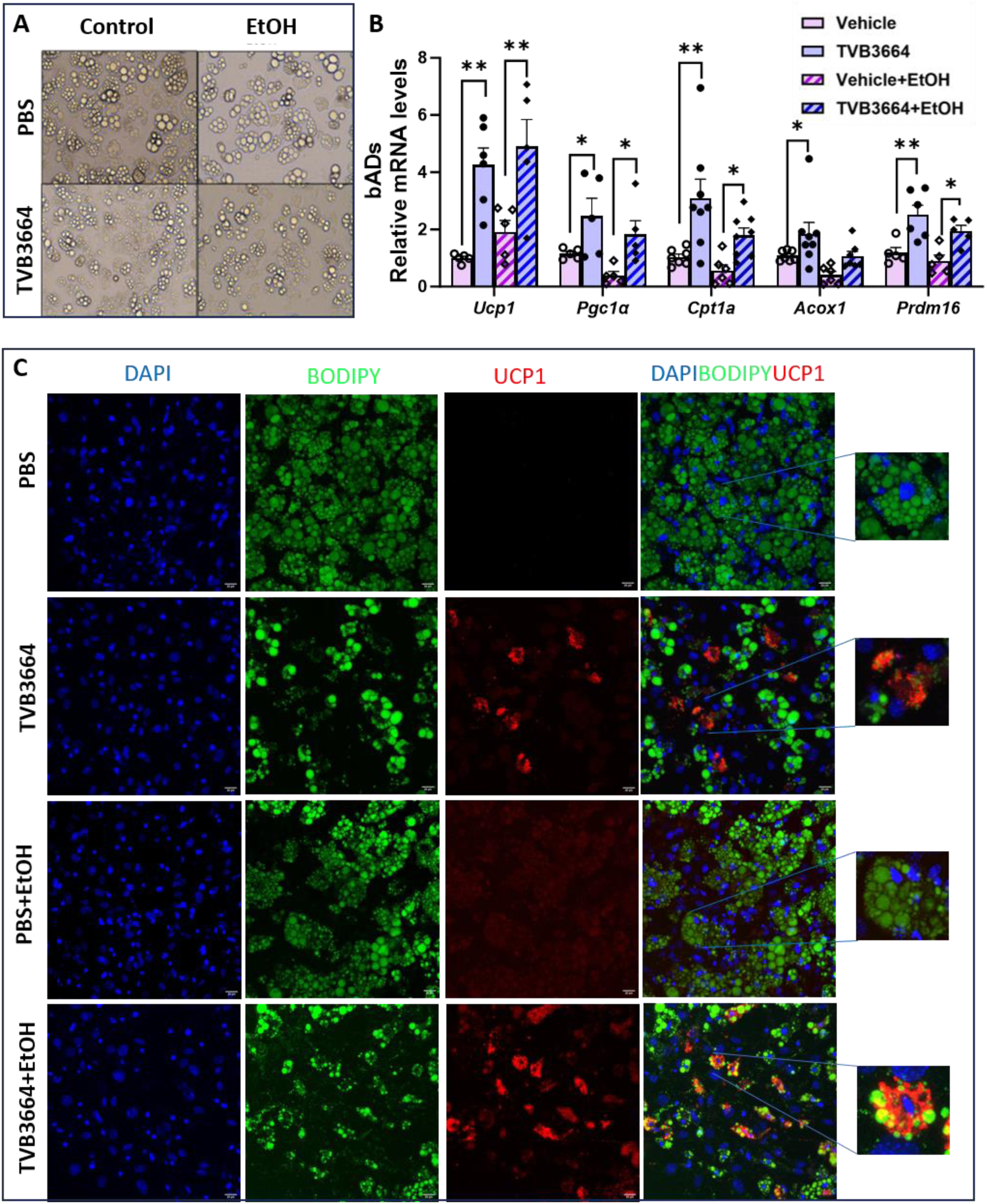
Pharmacological FASN inhibition in differentiated bADs induces thermogenic gene expression. Differentiated bADS were treated with TVB3664 or vehicle in the presence or absence of 100μM EtOH. (A) Representative images of differentiated bADs. (B) qPCR analysis of Ucp1, Pgc1α, Cpt1a, Acox1 and Prdm16 in bADS (n= 5-7). (C) Immunofluorescent staining of UCP1 and BODIPY in bADS. Data are expressed as the mean ± SEM. *P <.05, **P <.01.

### BAT-specific FASN deletion protects against acute alcohol-induced elevations in plasma TG in mice

Enhanced thermogenic program has been shown to promote plasma TG clearance and protect mice from hypertriglyceridemia ^34^. The elevated expression of UCP1 and other thermogenic genes in BAT of FASN^BKO^ mice led us to investigate the circulating TG levels in these mice. Interestingly, we found that acute alcohol exposure-induced increases in circulating TG was dramatically attenuated in FASN^BKO^ mice (**Figure 3A**). Bartlet et al. has demonstrated that cold exposure-induced enhancement of plasma TG clearance requires LPL and CD36 ^34^. However, FASN^BKO^ mice exhibited downregulated expressions of LPL and CD36 in both control and alcohol groups (**Figure 3B-C**). Recently, Shin et al. found that cold exposure induces VLDLR expression in BAT and VLDLR-mediated uptake of circulating VLDL particles contributes to lower plasma TG levels ^23^. To test this possibility, immunoblotting of VLDLR was performed and higher expression of VLDLR was observed in the BAT of acute alcohol-exposed FASN^BKO^ mice (**Figure 3C**). Then we performed VLDL-DiI uptake assay in differentiated bADs under both control and EtOH-exposed conditions. We found that regardless of the treatment, dramatically increased VLDL-DiI particles were observed in TVB3664-treated mature bADs (**Figure 3D** and **Supplementary Figure 4**). Functionally, VLDL-derived lipids are utilized to support thermogenic activity ^23^. Indeed, in the presence of VLDL, further elevated expression of thermogenic genes (Ucp1, Prdm16, PPARα) was observed in TVB3664-treated bADs (**Figure 3E**). Consistent with previous reports^35^, VLDL supplementation in vehicle-treated bADs did not alter the expression of thermogenic genes (**Figure 3E**). Interestingly, higher VLDLR mRNA expression was observed in bADs following TVB3664 plus VLDL-Dil incubation (**Figure 3E**). Furthermore, Acox1 expression was elevated in TVB3664-treated bADs under both VLDL-treated and untreated conditions, indicating enhanced β-oxidation (**Figure 3E**). Collectively, these findings support a model in which BAT-specific FASN deletion protects mice from alcohol-induced increases in circulating TG through increased VLDL particle uptake in the BAT.

**Figure 3.**
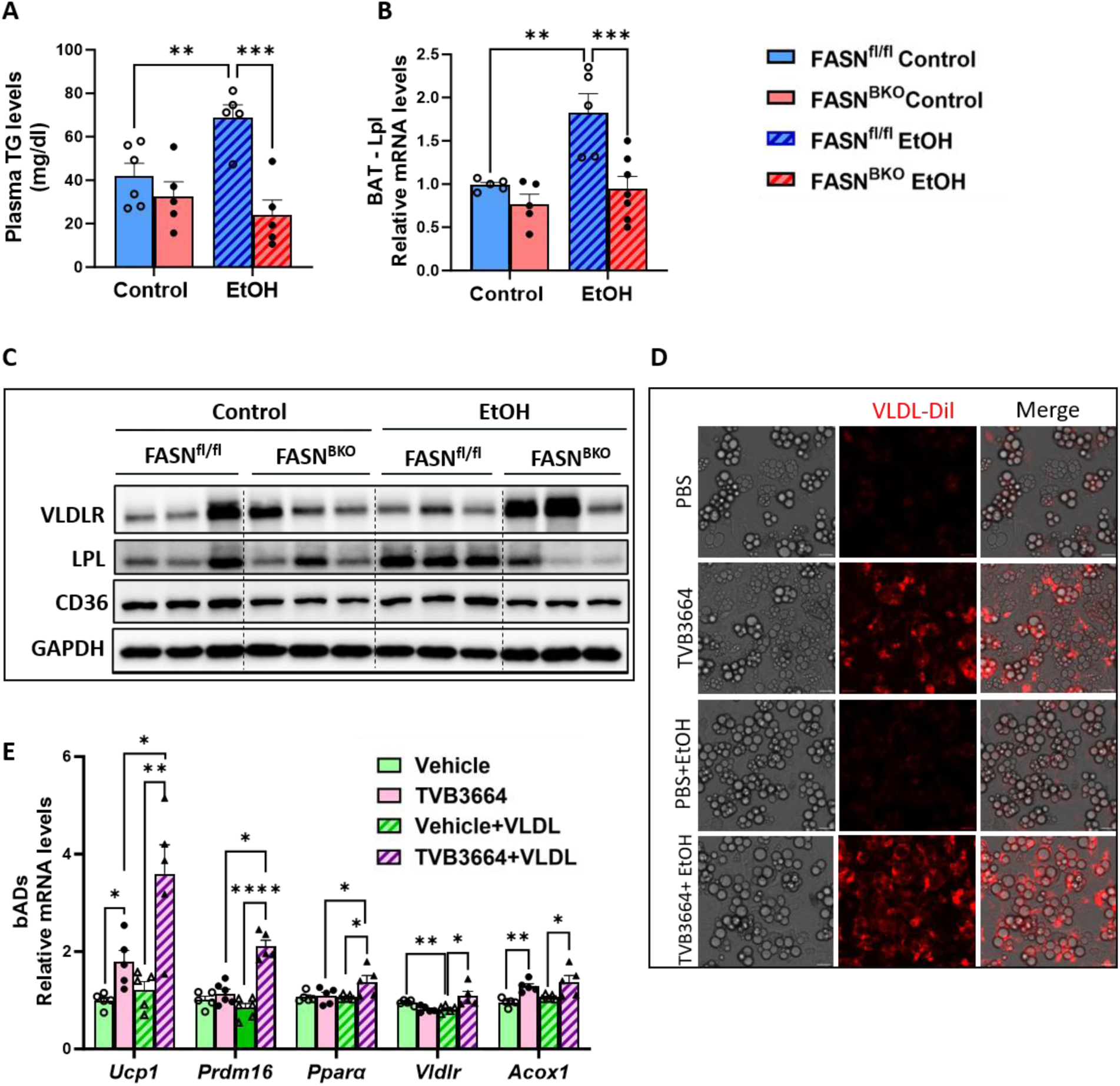
FASN inhibition in brown adipocytes promotes VLDL uptake and attenuates plasma TG levels. Male FASN^fl/fl^ and FASN^BKO^ mice were subjected to a single alcohol binge (EtOH, 5 g/kg BW) or isocaloric maltose dextrin (control) via oral gavage (A-C). (A) Plasma TG levels (n=5). (B) qPCR analysis of LPL in BAT (n=5-7). (C) Western blotting of VLDLR, LPL, CD36 and GAPDH in BAT. (D) Differentiated bADS were treated with TVB3664 or vehicle in the presence or absence of 100μM EtOH. Representative images of VLDL-Dil fluorescence in bADs. (E) Differentiated bADS were treated with TVB3664 or vehicle in the presence or absence of VLDL (10μg/ml). qPCR analysis of Ucp1, Prdm16, Pparα, Vldlr, and Acox1 in bADs (n=5). Data are expressed as the mean ± SEM. *P <.05, **P <.01, ***P <.001, **** P<.0001.

### BAT-specific FASN deletion protects against acute alcohol-induced hepatic steatosis

Following binge drinking, markedly increased plasma ALT and hepatic TG content levels were observed in FASN^fl/fl^ mice (**Figure 4A-B**). In contrast, these parameters were significantly attenuated in acute alcohol-exposed FASN^BKO^ mice (**Figure 4A-B**). Histological analyses further supported these findings, as Oil Red O and H&E staining revealed pronounced lipid droplet accumulation in hepatocytes of alcohol-exposed FASN^fl/fl^ mice, which was substantially reduced in FASN^BKO^ mice (**Figure 4C-D**). To investigate the underlying mechanisms that contribute to less lipid accumulation in binge drinking-treated FASN^BKO^ mice, we examined hepatic expression of CD36, a fatty acid translocase responsible for transporting FFAs into cells^36^. Surprisingly, we observed elevated CD36 expression in FASN^BKO^ mice (**Figure 4E**). Then we determined the expression of genes involved in fatty acid oxidation and found significantly enhanced protein expression of sirtuin 1 (SIRT1) and cytochrome c oxidase subunit 4 (CoxIV) in both control-and alcohol-fed FASN^BKO^ mice (**Figure 4F**). In addition, higher liver mRNA expression of Cpt1a, Acox1, and Pgc1α was observed in alcohol-exposed FASN^BKO^ mice compared to FASN^fl/fl^ mice (**Figure 4G**). UCP1-Cre-meidated BAT FASN deletion did not affect liver FASN expression (**Figure 4H**). Despite elevated FASN mRNA expression in response to acute alcohol challenge in both genotypes (**Figure 4I**), similar protein levels of FASN were found regardless of treatment or genotype (**Figure 4H**). Srebp1c is a critical transcription factor that controls lipogenesis and SREBP1c global knockout mice are protected from alcohol-induced fatty liver disease ^37^. Significantly reduced mRNA expression of Srebp1c was observed in acute alcohol-exposed FASN^BKO^ mice compared to FASN^fl/fl^ littermates under the same treatment (**Figure 4J**), although no significant differences were observed in the protein expression of SCD1, a target gene of SREBP1c (**Figure 4H**). Chronic alcohol feeding has been shown to enhance VLDL secretion and contribute to higher plasma TG content ^17^. Similarly, we found that acute alcohol drinking also promoted hepatic VLDL production in FASN^fl/fl^ mice (**Figure 4K**). Interestingly, VLDL secretion was significantly reduced in binge alcohol-treated FASN^BKO^ mice, which was comparable to control-fed mice (**Figure 4K**). Taken together, these observations indicate that BAT-specific FASN deletion protects against acute alcohol-induced hepatic steatosis and liver injury, primarily through enhanced hepatic β-oxidation.

**Figure 4.**
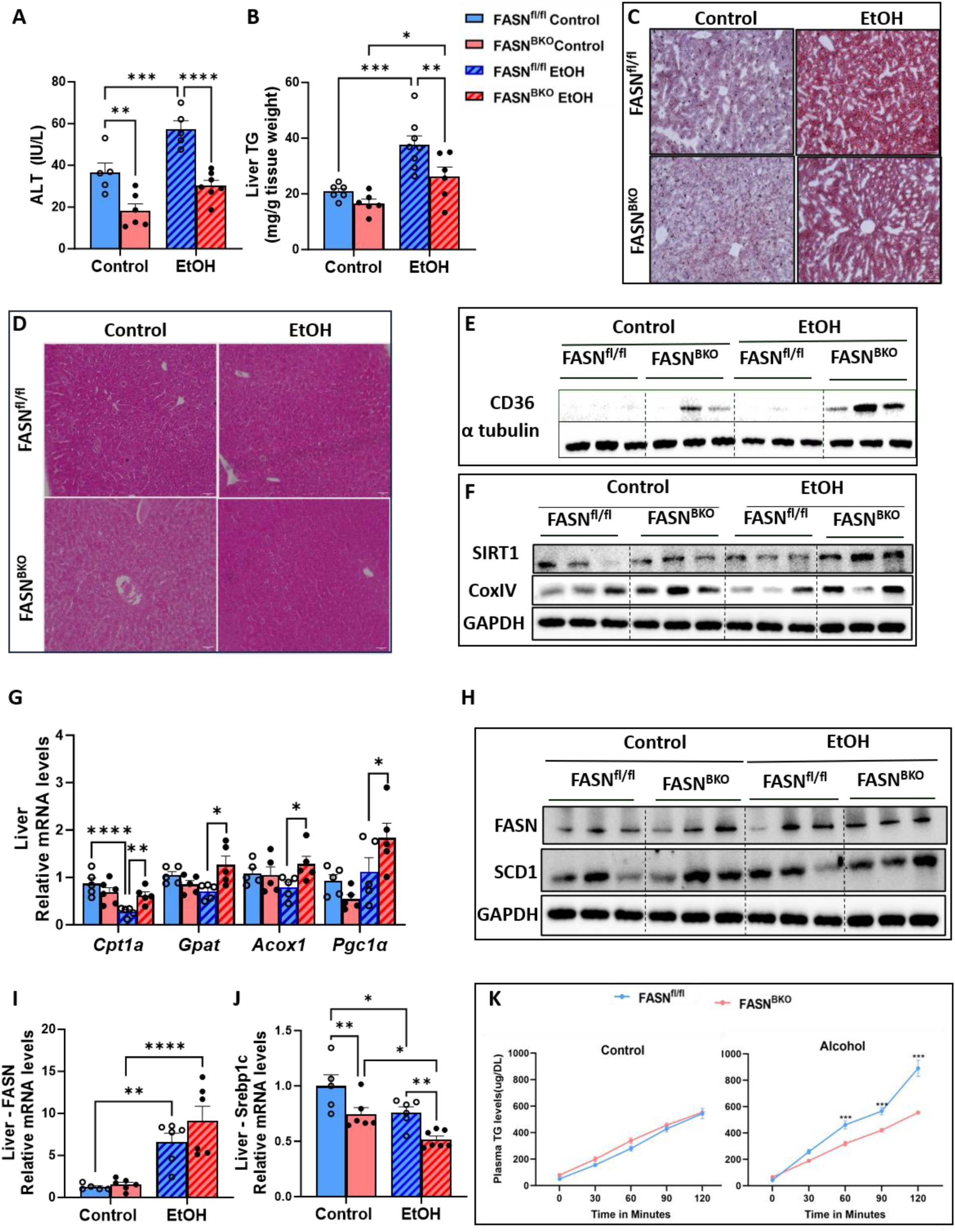
FASN inhibition in BAT rescues acute alcohol-induced liver lipid accumulation. Male FASN^fl/fl^ and FASN^BKO^ mice were subjected to a single alcohol binge (EtOH, 5 g/kg BW) or isocaloric maltose dextrin (control) via oral gavage. (A) Plasma ALT levels (n=5). (B) Liver TG levels (n=6-8). (C) Representative images of Oil Red O staining of liver tissues. (D) Representative images of H&E staining in liver tissues. (E and F) Western blotting of CD36, α tubulin, SITR1, CoxIV, and GAPDH in liver tissues. (G) qPCR analysis of Cpt1a, Gpat, Acox1 and Pgc1α in livers (n=5-6). (H) Western blotting of FASN, SCD1 and GAPDH in the liver. (I-J) qPCR analysis of liver FASN and Srebp1c (n= 5-7). (K) Hepatic VLDL production assay showing plasma TG levels at different time points (n=5). Data are expressed as the mean ± SEM. *P <.05, **P <.01, ***P <.001, **** P<.0001.

### BAT-specific FASN deletion enhances thermogenesis and protects against alcohol-induced increases in plasma TG and hepatic steatosis in mice following acute-on-chronic alcohol feeding

Acute-on-chronic alcohol exposure paradigm has been widely used to mimic the progression of alcohol-induced liver injury observed in humans. Prior reports have shown that this feeding regimen causes elevated UCP1 expression in the BAT ^38^. Consistently, significantly elevated mRNA levels of Ucp1, Pgc1a and Prdm16 as well as UCP1 protein expression were observed in alcohol-fed FASN^fl/fl^ mice (**Figure 5A-B**). Notably, the expression of these thermogenic genes was further elevated in the BAT of acute-on-chronic alcohol-exposed FASN^BKO^ mice (**Figure 5A-B**). In contrast to the similar expression of CPT1a and TOM20 in the BAT of control and acute alcohol-treated FASN^fl/fl^ mice (**Figure 1F**), following acute-on-chronic alcohol treatment, the expression of these two proteins was greatly increased in FASN^fl/fl^ mice (**Figure 5C**). Regardless of the treatment, BAT FASN deficiency caused increases in CPT1a and TOM20 expression, which were comparable to that of observed in alcohol-fed FASN^fl/fl^ mice. Despite higher BAT/body weight ratio in alcohol-exposed FASN^BKO^ mice (**Figure 5D**), these mice exhibited slightly reduced Cidea expression and greatly decreased lipid accumulation in the BAT (**Figure 5A and 5E**).

**Figure 5.**
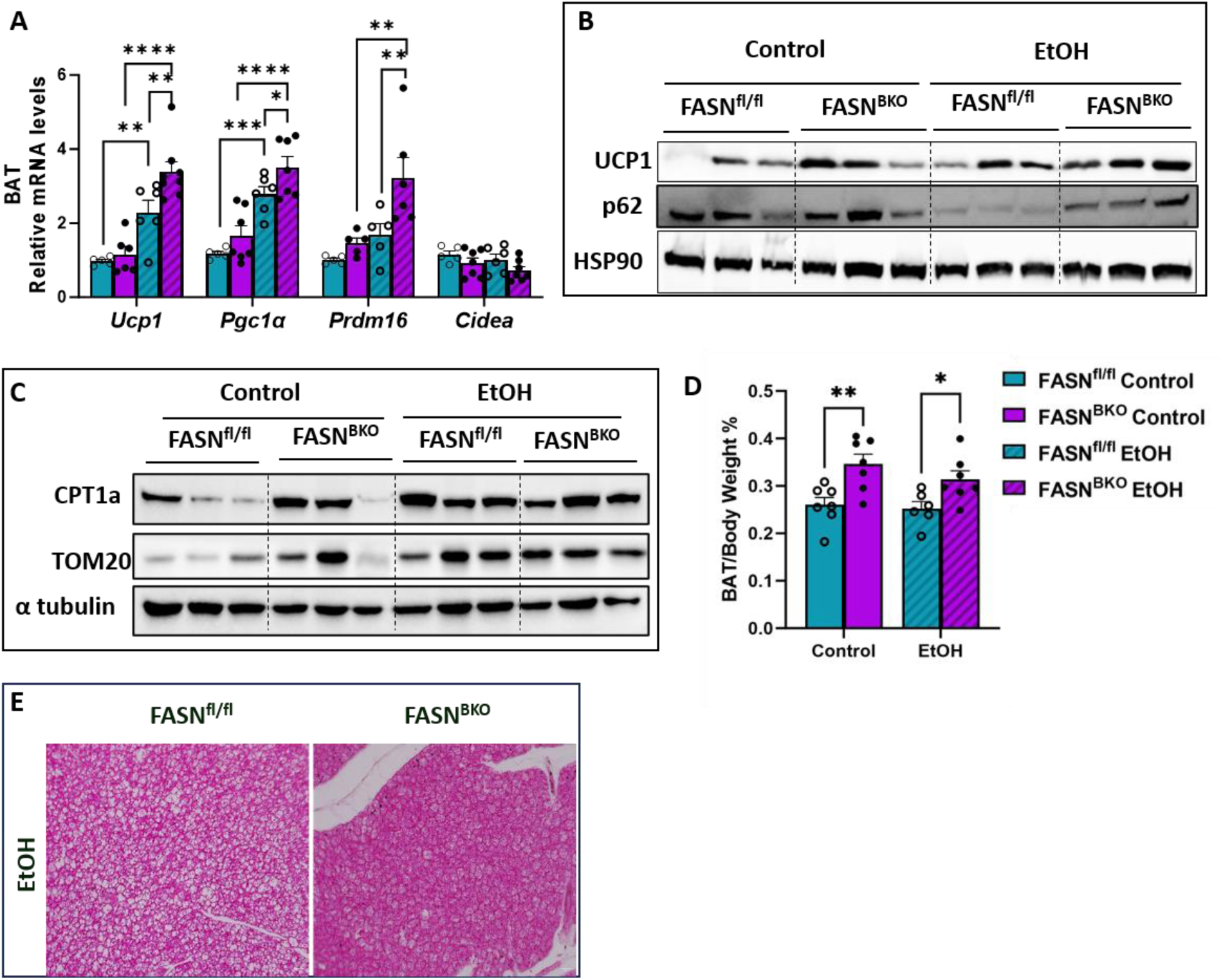
BAT-specific FASN deletion enhances thermogenesis in mice following acute-on-chronic alcohol feeding. Male FASN^fl/fl^ and FASN^BKO^ mice were subjected to 10 days of 5% alcohol diet or isocaloric maltose dextrin treatment. On the 11^th^ day, mice were administered a single oral gavage of alcohol (EtOH; 5 g/kg body weight) or an isocaloric maltose dextrin solution (control). (A) qPCR analysis of Ucp1, Pgc1α, Prdm16, and Cidea in BAT. (B) Western blotting of UCP1, p62 and HSP90 in BAT. (C) Western blotting of CPT1a, TOM20 and α tubulin in BAT. (D) BAT tissue weight/Body weight % (n=6-7). (E) Representative images of H&E staining in BAT. Data are expressed as the means ± SEM. *P <.05, **P <.01, ***P <.001, **** P<.0001.

Next, we set out to determine whether an enhanced thermogenic program in BAT of mice following acute-on-chronic alcohol exposure is correlated with reduced plasma TG. Similar to our observations in binge drinking-exposed mice, acute-on-chronic alcohol caused a great elevation in plasma TG in FASN^fl/fl^ mice, which were significantly attenuated in FASN^BKO^ mice (**Figure 6A**). Then we performed ex vivo VLDL-Dil uptake assay in mouse BAT as previously described ^23^. We found that FASN^fl/fl^ mice showed an increase in VLDL-Dil uptake following alcohol exposure and FASN^BKO^ mice exhibited markedly higher VLDL-Dil uptake under both control and alcohol conditions (**Figure 6B**). Acute-on-chronic alcohol feeding caused increases in liver weight to body weight ratio, hepatic TG content, and plasma ALT levels in FASN^fl/fl^ mice (**Figure 6C-E**). In contrast, these alcohol-induced liver injury markers were significantly attenuated in FASN^BKO^ mice (**Figure 6C-E**). Moreover, H&E staining demonstrated reduced hepatic lipid droplet accumulation in alcohol-fed FASN^BKO^ mice (**Figure 6F**). Mechanistically, these findings were associated with a significant induction of hepatic β-oxidation, as evidenced by markedly elevated mRNA and protein levels of CPT1a (**Figure 6G-H**) as well as increased protein expression of CoxIV and cytochrome c in alcohol-fed FASN^BKO^ mice (**Figure 6H**). Interestingly, in response to acute-on-chronic alcohol feeding, FASN^fl/fl^ mice exhibited decreased Cpt1a mRNA expression but increased CPT1a protein levels (**Figure 6G–H**). In comparison, FASN^BKO^ mice had increased Cpt1a mRNA and CPT1a protein levels following alcohol exposure (**Figure 6G–H**). It has been reported that mice subjected to acute-on-chronic alcohol feeding exhibit increased expression of Srebp1c and its target lipogenic genes such as FASN and SCD1 ^17^. Similar findings were observed in alcohol-fed FASN^fl/fl^ mice despite a slight reduction in FASN expression after alcohol treatment (**Figure 6I-J**). Interestingly, livers from alcohol-exposed FASN^BKO^ mice displayed marked downregulation of Srebp1c and SCD1, despite no significant change in FASN expression (**Figure 6I-J**). Previous studies reported that chronic alcohol feeding downregulates hepatic mRNA expression of Gpat1, a gene involved in liver TG metabolism ^39^. Similar findings were observed in the liver of acute-on-chronic alcohol-fed FASN^fl/fl^ mice whereas Gpat1 expression remained largely unaffected in FASN^BKO^ mice (**Figure 6K**). Together, these findings demonstrate that inhibition of FASN in BAT promotes thermogenic activity and protects against elevations in plasma TG and hepatic fat accumulation in mice exposed to acute-on-chronic alcohol feeding.

**Figure 6.**
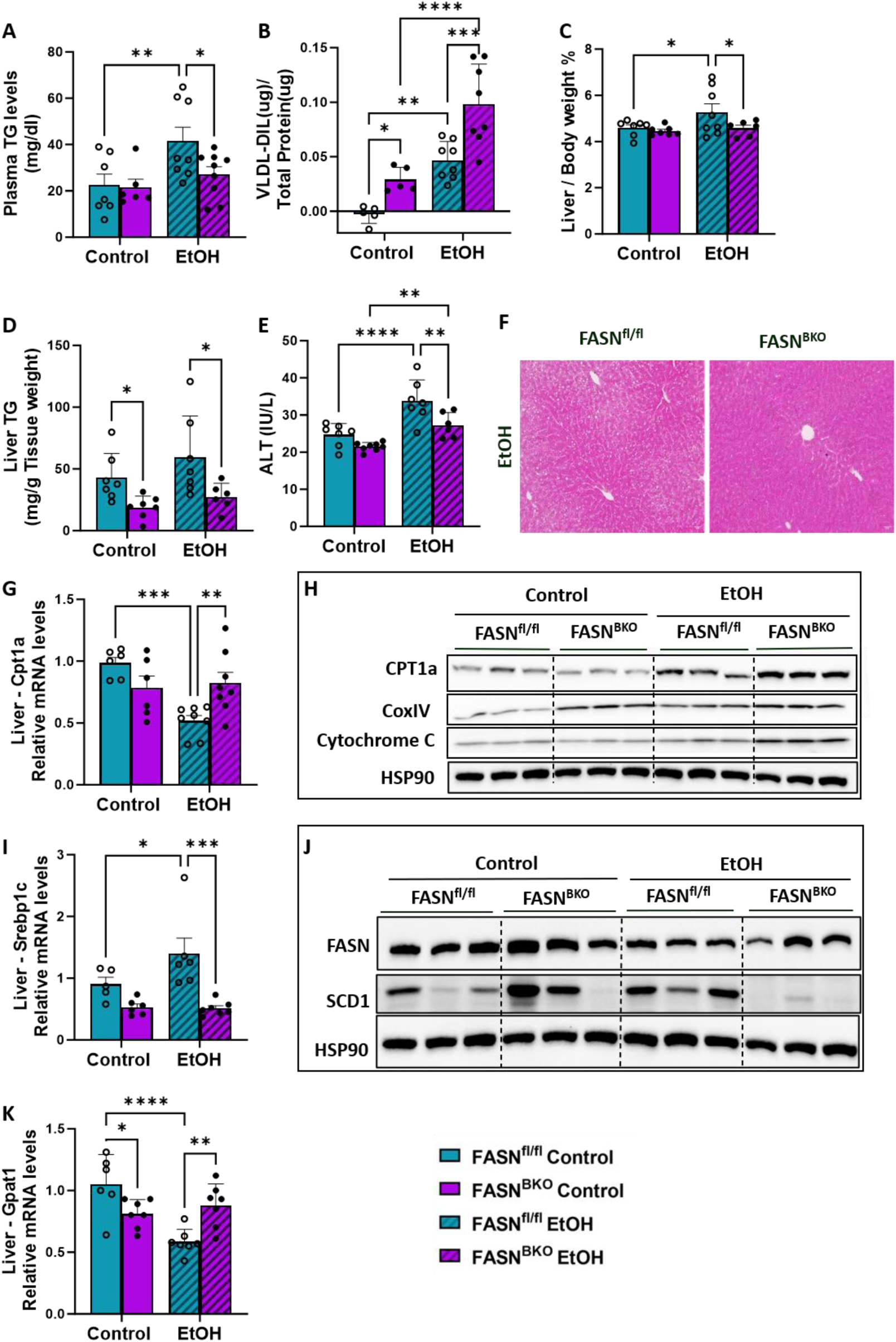
Bat-specific FASN deficiency attenuates acute-on-chronic alcohol-induced hepatic steatosis. Male FASN^fl/fl^ and FASN^BKO^ mice were subjected to 10 days of 5% alcohol diet or isocaloric maltose dextrin followed by a single oral gavage of alcohol (EtOH, 5 g/kg BW) or isocaloric maltose dextrin (control) on the 11^th^ day. (A) Plasma TG levels (n=6-8). (B) VLDL-Dil content in BAT (n=5-8). (C) liver weight/body weight % (n=7-8). (D) Liver TG levels (n=6-8). (E) Plasma ALT levels (n=5). (F) Representative images of H&E staining in liver tissues. (G) qPCR analysis of hepatic Cpt1a expression (n=6-8). (H) Western blotting of CPT1a, CoxIV, Cytochrome C and HSP90 in liver. (I) qPCR analysis of hepatic Srebp1c expression (n= 6-8). (J) Western blotting of FASN, SCD1 and HSP90 in liver. (K) qPCR analysis of hepatic Gpat1 expression (n=5-7). Data are expressed as the means ± SEM. *P <.05, **P <.01, ***P <.001, **** P<.0001.

### FASN^BKO^ mice show differential levels of circulating adipokines and inflammatory markers

To investigate the mechanisms through which FASN^BKO^ mice are protected from alcohol-induced liver injury, we performed Proteome Profiler Mouse Adipokine Antibody Arrays using plasma collected from acute-on-chronic alcohol-treated mice. Among the 38 adipokines analyzed, six proteins exhibited marked differences in FASN^BKO^ mice compared with FASN^fl/fl^ mice under both control and alcohol-fed conditions (**Figure 7A**).

**Figure 7.**
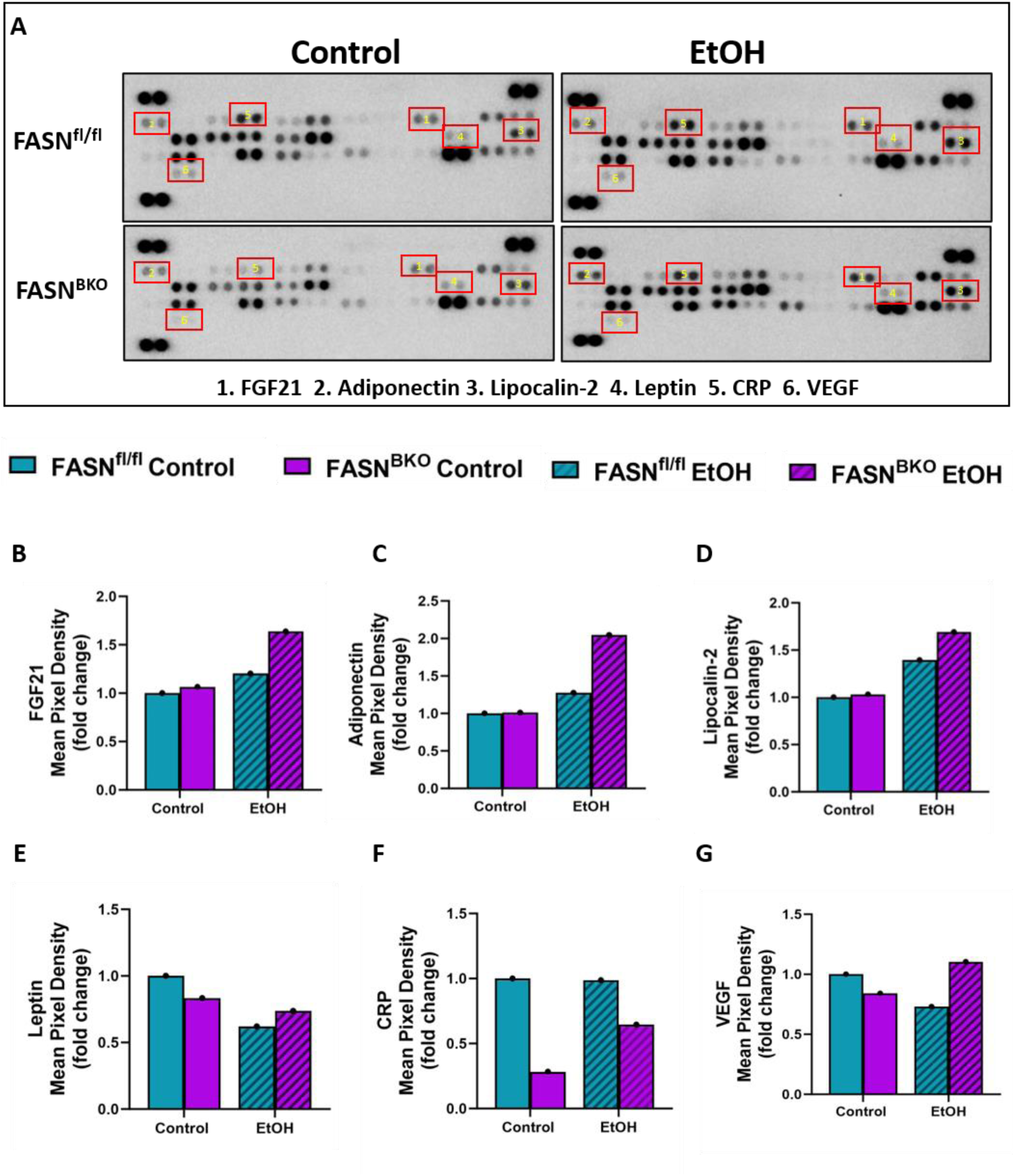
FASN^BKO^ mice show differential expression of adipokine and inflammatory markers in the circulation. Male FASN^fl/fl^ and FASN^BKO^ mice were subjected to 10 days of 5% alcohol diet or isocaloric maltose dextrin followed by a single oral gavage of alcohol (EtOH, 5 g/kg BW) or isocaloric maltose dextrin (control) on the 11^th^ day. (A) Images of plasma adipokine array. Plasma from 8 mice were pooled for the assay. (B-G) Densitometry analysis of selected adipokines including FGF21, adiponectin, Lipocalin-2, Leptin, CRP, and VEGF from the adipokine array.

Plasma levels of fibroblast growth factor 21 (FGF21), adiponectin, and lipocalin-2 demonstrated substantial changes in FASN^BKO^ mice specifically after alcohol exposure (**Figure 7B-D**). These adipokines are known to play important roles in regulating systemic metabolism, inflammation, and stress responses ^40–42^. Leptin levels were reduced in FASN^BKO^ mice under control conditions, whereas plasma leptin levels were increased in alcohol-exposed FASN^BKO^ mice. (**Figure 7E**). Notably, plasma levels of C-reactive protein (CRP), a well-established marker of systemic inflammation, were greatly reduced in alcohol-fed FASN^BKO^ mice (**Figure 7F**). Consistent with this reduction, the inflammatory markers (IL-1β and MCP1) were significantly reduced in the liver of acute-on-chronic alcohol-fed FASN^BKO^ mice (**Supplementary Figure 5**). These findings suggest an overall attenuation of inflammatory responses in FASN^BKO^ mice and may contribute to their resistance to alcohol-induced liver injury. In addition, BAT thermogenic activation has previously been associated with increased secretion of vascular endothelial growth factor (VEGF), which promotes angiogenesis and vascular remodeling ^43–45^. Consistent with enhanced thermogenic activity, we observed a modest increase in plasma VEGF levels in alcohol-fed FASN^BKO^ mice (**Figure 7G**). Overall, these findings suggest that FASN^BKO^ mice exhibit a distinct circulating adipokine profile and these alterations may represent important mechanisms through which increased BAT thermogenesis confers protection against alcohol-induced liver damage.

### BAT FASN deficiency is associated with activation of the FGF21-AMPK pathway

BAT-derived FGF21 has been proposed to function as a batokine that contributes to inter-organ metabolic communication ^46–48^. Based on the array data in **Figure 7**, we determined adiponectin and FGF21 expression in mouse BAT. Consistent with array data in **Figure 7C**, plasma adiponectin levels were higher in FASN^BKO^ mice (**Supplementary Figure 6A**). However, FASN deficiency or alcohol treatment did not affect adiponectin mRNA expression in BAT and in bADs (**Supplementary 6B-D**). Interestingly, FASN^BKO^ mice exhibited significantly increased FGF21 expression in BAT compared with FASN^fl/fl^ littermates under both control and alcohol conditions in both acute and acute-on-chronic treatment paradigms (**Figure 8A-B**). Notably, circulating FGF21 levels were also elevated in FASN^BKO^ mice following control and acute alcohol treatment and alcohol-fed FASN^BKO^ mice displayed the highest levels of FGF21 (**Figure 8C**). For mice exposed to acute-on-chronic feeding regimen, FASN^BKO^ mice exhibited significantly higher plasma FGF21 levels after control diet exposure (**Figure 8D**). However, slightly but not significantly higher circulating FGF21 content was observed in alcohol-fed FASN^BKO^ mice (**Figure 8D**). To determine whether FASN deficiency-mediated FGF21 regulation may occur intrinsically, we further assessed FGF21 expression in FASN-inhibited bADs in vitro and found significantly increased FGF21 mRNA expression in TVB3664-treated bADs (**Figure 8E**). In addition, greatly elevated levels of FGF21 were observed in the culture medium collected from TVB3664-treated bADs following both control and alcohol treatments (**Figure 8F**). Zhu et al. showed that the recombinant FGF21 treatment ameliorates the development of alcoholic fatty liver in mice via FGFR1 and pAMPK mediated signaling ^49^. Figure 8G and 8H showed elevated hepatic mRNA expression of FGFR1 in alcohol-fed FASN^BKO^ mice. In addition, significantly increased hepatic pAMPK levels were observed in FASN^BKO^ mice under both acute and acute-on-chronic alcohol conditions (**Figure 8I-N**). Collectively, these findings indicate an association between BAT FASN deletion and elevation of the FGF21 levels and suggest that BAT-derived FGF21 may contribute to the improved hepatic metabolic adaptations via activation of pAMPK pathway in alcohol-exposed FASN^BKO^ mice.

**Figure 8.**
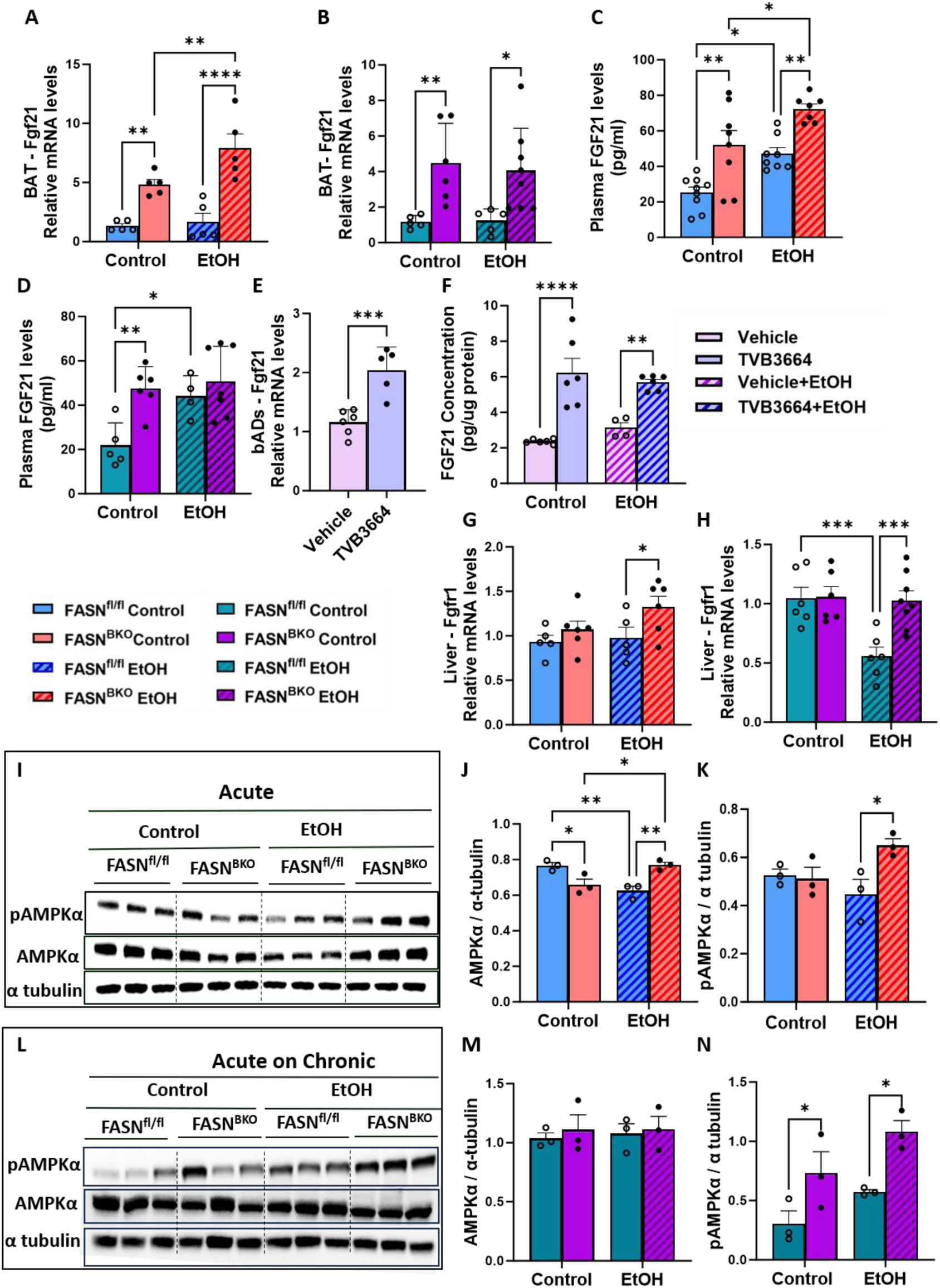
BAT FASN deficiency is associated with increased expression and production of FGF21. (A-B) qPCR analysis of FGF21 expression in the BAT of binge (A, n=5) and acute-on-chronic alcohol-fed mice (B, n=5-7). (C-D) Plasma FGF21 levels in binge (C, n= 8-11) and acute-on-chronic alcohol-fed mice (D, n= 5-7). (E) qPCR analysis of FGF21 expression in bADs (n=5-7). (F) FGF21 levels in culture media of differentiated bADs (n=4-6). (G-H) qPCR analysis of hepatic FGFR1 expression of binge (G, n=5-6) and acute-on-chronic alcohol-fed mice (H, n=6-8). (I) Western blotting of pAMPKα, AMPKα and α tubulin in livers of acute-alcohol-fed mice. (J-K) Densitometry analysis of AMPKα (J) and pAMPKα (K) from I. (L) Western blotting of pAMPKα, AMPKα and α tubulin in livers of acute-on-chronic alcohol-fed mice. (M-N) Densitometry analysis of AMPKα (M) and pAMPKα (N) from L. Data are expressed as the mean ± SEM. *P <.05, **P <.01, ***P <.001.

## Discussion

Consistent with a recent report that BAT-specific FASN deficiency leads to elevated UCP1 expression and reduced lipid droplet accumulation in the BAT of chow-fed male mice, we found that, regardless of diet treatment, FASN^BKO^ mice exhibited upregulated expression of UCP1 and multiple thermogenic genes as well as less lipid accumulation within BAT. Both studies have used UCP1-Cre (JAX:024670) to mediate FASN ablation. Given the findings from recent lineage-tracing studies showing that constitutive UCP1-Cre activity is not completely restricted to BAT but can be detected in non-adipose tissues, including regions of the central nervous system, kidney, adrenal gland, and other developmental lineages ^50,51^, we isolated and differentiated bADs to investigate the effect of FASN deficiency on UCP1 expression. Importantly, our in vitro assay showed that TVB3664-mediated inhibition of FASN in differentiated bADs reproduced the major metabolic adaptations observed in vivo, including enhanced thermogenesis and reduced lipid droplet accumulation. These complementary genetic and pharmacological approaches strongly support the direct role of brown adipocyte FASN in regulating thermogenic function.

Rowland et al. reported that adipocyte-specific deletion of FASN resulted in increased p62 levels in various adipose tissues, including BAT ^28^. In line with these findings, our study demonstrated that BAT-specific FASN ablation increased p62 protein levels, a phenotype that was further corroborated in differentiated bADs following pharmacological inhibition of FASN with TVB3664. Accumulating evidence has shown that p62 functions as an important regulator of bADs thermogenesis. Fischer et al. found that BAT-specific p62 deficient mice have markedly reduced UCP1 expression in BAT and overexpression of p62 in p62-deficient primary bADs restores thermogenic gene expression ^52^. Additionally, Qian et al. reported that aged p62 knockout mice display impaired BAT function and develop more severe liver injury upon alcohol exposure ^53^. Collectively, these findings raise the possibility that elevated p62 levels observed in FASN ^BKO^ mice and TVB3664-treated bADs may partially contribute to the enhanced UCP1 expression and thermogenic phenotype observed in our models.

The most striking consequence of BAT FASN deletion was the protection against alcohol-induced elevations in plasma TG. Activated BAT is recognized as an important site for clearance of circulating triglyceride-rich lipoproteins through LPL-CD36 mediated uptake and oxidation of lipid-derived substrates ^34^. However, alcohol-fed FASN^BKO^ mice displayed reduced expression of LPL and CD36. Interestingly, increased expression of VLDLR was observed in the BAT of FASN^BKO^ mice under both acute and chronic alcohol conditions. Shin et al. reported that VLDLR expression on bADs and BAT mediates VLDL uptake and contributes to enhanced plasma TG clearance in cold-induced thermogenesis^23^. Consistent with these findings, markedly increased VLDL-Dil particle uptake was observed in the BAT of FASN^BKO^ mice and in the differentiated bADs treated with FASN inhibitor TVB3664, supporting a role of BAT FASN inhibition in promoting VLDL-derived lipid acquisition and lowering plasma TG. This increased utilization of VLDL-derived fatty acids may additionally provide substrates required to sustain thermogenic activity. Indeed, Shin et al. demonstrated that following uptake into BAT, VLDL particles are processed through lysosomal pathways, resulting in the release of FFAs that can be utilized for thermogenesis ^23^. Thus, BAT FASN deletion appears to rescue hypertriglyceridemia by clearing VLDL from the circulation and utilizing it as a source for thermogenesis.

Several studies have demonstrated that impaired thermogenic activity contributes to increased hepatic steatosis during chronic and acute-on-chronic alcohol feeding models. Shen et al. showed that whole-body UCP1 deficiency exacerbates alcohol-induced hepatic steatosis in both male and female mice, highlighting the protective role of thermogenesis in maintaining hepatic lipid homeostasis ^30^. Similarly, BAT-specific deletion of G protein-coupled bile acid receptor 1 (TGR5) results in suppressed thermogenic activity and increased liver steatosis under alcohol exposure conditions ^29^. Conversely, pharmacological activation of thermogenesis through a TGR5 agonist is able to alleviate alcohol-induced hepatic steatosis ^29^. In agreement with these findings, our study showed that FASN^BKO^ mice displayed upregulated expression of thermogenic genes in BAT and are protected from hepatic steatosis induced by both binge drinking and acute-on-chronic alcohol feeding. Furthermore, we found that this hepatic protection observed in alcohol-fed FASN^BKO^ mice was associated with improved mitochondrial oxidative metabolism, indicated by increased expression of mitochondrial proteins including COXIV and cytochrome c and genes involved hepatic fatty acid oxidation (Cpt1a, Acox1, Pgc1α).

FGF21 is predominantly produced and secreted by the liver, where fasting and metabolic stress induce hepatic FGF21 expression through PPARα signaling^54^. Liver-derived FGF21 acts as an endocrine hormone to regulate systemic energy metabolism, including fatty acid oxidation, ketogenesis, and lipid homeostasis^55–58^. Interestingly, recent studies have revealed that FGF21 can also be produced by extrahepatic tissues, including skeletal muscle and BAT ^59–61^. Specifically, Hondares et al. demonstrated that thermogenic activation induces FGF21 expression and secretion from rodent BAT and cultured differentiated bADs, establishing BAT as an endocrine source of FGF21 ^60^. In addition, Pereira et al. showed that BAT mitochondrial stress stimulates activating transcription factor 4 (ATF4)-dependent FGF21 production from BAT and cultured bADs^59^. In the current study, both proteome array and ELISA analysis revealed a striking increase in circulating FGF21 levels in both control-and alcohol-fed FASN^BKO^ mice. Notably, TVB3664-mediated inhibition of FASN in cultured bADs resulted in increased FGF21 mRNA expression and enhanced secretion of FGF21 into the culture medium. These observations indicate that FASN inhibition could promote FGF21 production in bADs and BAT.

Several studies have demonstrated that FGF21 plays a pivotal role in maintaining hepatic lipid homeostasis through the activation of AMPK-mediated signaling pathway ^62–64^. Yano et al. found that hepatocyte-specific FGF21 overexpression enhances the phosphorylation of AMPK and promotes hepatic fatty acid oxidation, contributing to attenuated liver steatosis in high-fat diet-fed mice^65^. Moreover, Zhu et al. reported that FGF21 administration activates the AMPK-SIRT1 pathway and attenuates lipid accumulation in livers of alcohol-fed mice and in alcohol-exposed HepG2 cells^66^. Consistently, we observed increased hepatic pAMPK levels in FASN^BKO^ mice following both binge drinking and acute-on-chronic alcohol exposure, which was accompanied by elevated SIRT1 protein expression and protection against alcohol-induced hepatic steatosis.

The observed increase in VLDLR expression in the BAT of FASN^BKO^ mice may represent a compensatory adaptation to the loss of de novo fatty acid synthesis. BAT relies on multiple lipid sources to sustain thermogenesis, including intracellular triglyceride stores, de novo lipogenesis, and the uptake of circulating triglyceride-rich lipoproteins^67^. In the absence of brown adipocyte FASN, the reduced capacity for endogenous fatty acid synthesis is likely to increase the dependence of BAT on exogenous lipid uptake to meet the high energetic demands of thermogenesis. Previous studies have demonstrated that VLDLR is a stress-responsive receptor whose expression is induced under conditions of metabolic stress. Yang et al. showed that hypoxia induces VLDLR expression in cultured endothelial cells as part of a stress response associated with endoplasmic reticulum stress and apoptosis^68,69^. Using primary hepatocytes as well as liver samples from mice and humans, Zarei et al. reported that hepatic VLDLR is regulated by PPARβ/δ-FGF21 signaling axis during metabolic adaptation in non-alcoholic fatty liver disease^70^. In the current study, FASN deficiency in BAT creates a metabolic state characterized by reduced endogenous lipid availability due to impaired de novo lipogenesis and increased substrate demand evidenced by enhanced expression of thermogenic genes. This metabolic adaptation may trigger VLDLR expression to facilitate the uptake of exogenous lipids. Future studies are warranted to determine the molecular mechanisms linking FASN deficiency to VLDLR induction in BAT.

In summary, BAT-specific FASN deficiency promotes the clearance of triglyceride-rich VLDL particles during alcohol exposure, thereby lowering plasma TG levels in mice. In addition, BAT-specific FASN deletion attenuates alcohol-induced hepatic steatosis, likely through BAT-liver metabolic crosstalk.

## Materials and Methods

### Animals

C57BL/6J mice were obtained from the Jackson laboratory. The generation and validation of FASN^fl/fl^ mice have been previously described^71^. To selectively delete FASN in BAT, UCP1-Cre transgenic mice (JAX:024670) were crossed with FASN^fl/fl^ mice to produce mice lacking FASN specifically in BAT (FASN^BKO^). Animals were housed in a pathogen-free barrier facility with a 12-hour light-dark cycle (6:00 a.m.-6:00 p.m.). Mice had free access to food (standard chow diet, 2916 Global Diet; Harlan Teklad) and water unless specified otherwise. Experiments were performed according to protocols reviewed and approved by the Institutional Animal Care and Use Committee of the University of Texas at Dallas (UTD).

### Alcohol Treatment and Tissue collection

Male FASN^fl/fl^ and FASN^BKO^ mice (8-9 weeks old, at least 20g in weight) were given an oral gavage of 5 g/kg BW of alcohol (31.5%, vol/vol, referred to as EtOH) or maltose dextrin (45%, wt/vol, referred to as control). Six hours later, mice were anesthetized for blood, liver and BAT collection. For acute-on-chronic treatment mice were given Lieber – Decarli ‘82 Diet 5% alcohol diet, or isocaloric control diet for 10days and on 11^th^ day mice were given one oral gavage similar to one binge. Six hours later, mice were anesthetized for blood, liver, and BAT collection. Tissues from both the treatments were quickly removed, snap-frozen in liquid nitrogen, and stored at −80^°^C.

### Cell Culture

Brown adipocyte-specific stromal vascular fraction (SVF) cells were isolated from BAT of C57BL/6J mice using a standard isolation protocol^72^. Briefly, dissected BAT tissues were minced and enzymatically digested in a digestion buffer containing collagenase D (Fisher Scientific, Cat# 17-101-015) and Dispase II (Sigma-Aldrich, Cat# D4693). Following digestion, the SVF fraction was separated through sequential centrifugation steps and filtration. Isolated SVF cells were cultured in DMEM/F-12 medium (Fisher Scientific, Cat# MT10090CV) supplemented with 15% fetal bovine serum (FBS). Upon reaching 100% confluence, cells were reseeded at a density of 0.5-1 × 10⁶ cells per dish and maintained in growth medium for 24 hours. Differentiation was initiated by replacing the growth medium with induction medium consisting of DMEM/F-12 supplemented with 10% FBS, 0.5mM 3-isobutyl-1-methylxanthine (IBMX; Sigma-Aldrich, Cat# I7018-100MG), 5μM dexamethasone (Sigma-Aldrich, Cat# D4902-25MG), 2μM rosiglitazone (Fisher Scientific, Cat# NC0950281), and 0.5μg/mL insulin (Sigma-Aldrich, Cat# I5500-50MG). After 48 hours of induction, cells were switched to a maintenance medium containing DMEM/F-12, 10% FBS, and 0.5μg/mL insulin. The maintenance medium was replaced every two days for 8 days until complete differentiation was achieved. For FASN inhibition, TVB3664 was used at a concentration of 100nM according to the previous report to effectively inhibit FASN activity in vitro^73^. Following 3 days treatment, conditioned media were collected and stored at −80°C for subsequent analysis. Cells were harvested for protein and RNA extraction.

### Histological Analysis

BAT and liver samples were fixed in 10% neutral-buffered formalin and processed for paraffin embedding by the Histology Core Facility at The University of Texas at Dallas (UTD). Paraffin-embedded liver and BAT tissues were sectioned at 5μm thickness and stained with hematoxylin and eosin (H&E). Whole-slide images were acquired using an EVOS M5000 Imaging System (Thermo Fisher Scientific). Some liver tissues were embedded in optimal cutting temperature (OCT) compound, cryo-sectioned at 5μm, and stained with Oil Red O according to standard procedures. Images were captured using the EVOS M5000 Imaging System under identical acquisition settings.

### Measurement of Liver and BAT TG Content

Hepatic TG content was measured as previously described^74^. Briefly, approximately 60– 80mg of liver tissue was minced, and lipids were extracted overnight at room temperature in a 2:1 (v/v) chloroform solution. Following centrifugation, 0.05% H₂SO₄ was added to facilitate phase separation, and the upper aqueous phase was carefully removed. A 100-μL aliquot of the lower organic phase was transferred to a clean tube and dried under a stream of nitrogen (N₂). The dried lipid extract was resuspended in 1% Triton X-100 in chloroform, after which the chloroform was evaporated under nitrogen. Subsequently, 500μL of deionized water was added, and samples were vortexed until the solution became clear. We employed the same method to isolate BAT TG, but we used 200ul of the lower organic phase. TG concentrations were determined using the Infinity™ TG Reagent (Thermo Fisher Scientific, Waltham, MA, USA) according to the manufacturer’s instructions.

### Western Blotting

Liver, BAT, and bADs samples were homogenized in radioimmunoprecipitation assay (RIPA) buffer containing 25mM Tris-HCl (pH 8.0), 150 mM NaCl, 1% NP-40, 1% sodium deoxycholate, and 0.1% SDS, supplemented with protease inhibitor cocktail (P8340, Sigma-Aldrich) and phosphatase inhibitor cocktails (P5726 and P0044, Sigma-Aldrich). Protein concentrations were determined using the Pierce™ BCA Protein Assay Kit (Thermo Fisher Scientific) according to the manufacturer’s instructions. Equal amounts of protein were resolved by SDS-PAGE and transferred onto nitrocellulose membranes (Trans-Blot®, Bio-Rad). Membranes were blocked with 5% nonfat dry milk or 3% bovine serum albumin (BSA) for phosphorylated proteins for 1 h at room temperature, followed by incubation with primary antibodies overnight at 4°C. Primary antibodies were diluted in the corresponding blocking buffer (5% nonfat dry milk or 3% BSA for phospho-specific antibodies). After washing with Tris-buffered saline containing 0.1% Tween-20 (TBST), membranes were incubated with horseradish peroxidase (HRP)-conjugated goat anti-rabbit IgG secondary antibody (Jackson Immuno Research) or anti-mouse IgG secondary antibody (1:10,000 dilution) for 1h at room temperature. Immunoreactive bands were detected using enhanced chemiluminescence (ECL) reagents and visualized with a chemiluminescence imaging system (Bio-Rad).

### Immunofluorescence staining

BAT samples were fixed in 4% paraformaldehyde (PFA), embedded in OCT compound, and cryo-sectioned for immunofluorescence staining. Tissue sections were permeabilized and blocked for 1 h at room temperature in blocking buffer (1× PBS containing 5% normal serum and 0.3% Triton X-100) before incubation with primary antibodies overnight at 4°C. After washing with PBS, sections were incubated with the appropriate fluorophore-conjugated secondary antibodies for 1h at room temperature. Nuclei were counterstained with 4′,6-diamidino-2-phenylindole (DAPI), and sections were mounted using an antifade mounting medium. Brown adipocytes (bADs) cultured on coverslips were fixed with 4% paraformaldehyde (PFA) and processed for immunofluorescence staining using the primary and secondary antibodies as described above. Fluorescence images were acquired using a fluorescence microscope (Olympus FV4000RS) under identical imaging settings for all experimental groups.

**Table 1:** List of antibodies.

| Antibody | Catalog# | Company |
| --- | --- | --- |
| UCP1 | 14670S | CST |
| GAPDH | 2118S | CST |
| $\alpha$ tubulin | 2125S | CST |
| FASN | 3180S | CST |
| SQSTM1/p62 (D6M5X) | 23214 | CST |
| UCP1(IF) | 72298S | CST |
| AMPKα | 2532S | CST |
| pAMPKα | 2535S | CST |
| CD36 | 74002 | CST |
| SCD1 | 2794S | CST |
| SIRT1 | 9475T | CST |
| Cpt1a | 12252T | CST |
| Cytochrome C | 4280S | CST |
| Cox IV | 4850S | CST |
| LPL | G2523 | Santa cruz |
| HSP90 | 4877T | CST |
| Tom20 | 42406S | CST |
| VLDLR | D138764-3 | NOVUS |
| HRP conjugated goat anti-mouse IgG | NC9491974 | Jackson Immuno Research |
| HRP conjugated goat anti-rabbit IgG | NC9611376 | Jackson Immuno Research |

### Plasma Parameters

Mice were anesthetized, and blood was collected into EDTA-coated tubes. Plasma was separated by centrifugation at 8,000 × *g* for 15 min and stored at −80°C until analysis. Plasma alanine aminotransferase (ALT) activity was measured using a commercial assay kit (Teco Diagnostics). Plasma fibroblast growth factor 21 (FGF21) levels were quantified using the Mouse/Rat FGF-21 Quantikine ELISA Kit (R&D Systems; Catalog No. MF2100). Plasma adiponectin levels were quantified using Mouse Adiponectin/Acrp30 Immunoassay (R&D Systems; Catalog No. MRP300, SMRP300 and PMRP300). Plasma triglyceride concentrations were measured using the Infinity™ TG Liquid Stable Reagent (Thermo Fisher Scientific; Catalog No. TR22421) according to the manufacturer’s protocol.

### VLDL Secretion Assay

Following a 4h fasting period after oral gavage of acute control or alcohol, mice were anesthetized and administered Tyloxapol (Triton WR-1339; Sigma) at a dose of 500 mg/kg body weight via retro-orbital vein injection to inhibit intravascular lipolysis. Blood samples were collected at 0, 30, 60, 90, and 120 min after Tyloxapol administration. Plasma TG concentrations were subsequently determined using an enzymatic assay as mentioned above.

### Cellular VLDL Uptake Assay

Following adipocyte differentiation and treatment with the FASN inhibitor (0.1µM working concentration) and alcohol (35µM or100µM), cellular VLDL uptake was assessed as previously described ^23^. Briefly, differentiated adipocytes were incubated with fluorescently labeled VLDL (VLDL-Dil, 10μg/ml) or unlabeled VLDL for 1 hour. Following incubation, cells were washed extensively with phosphate-buffered saline (PBS) to remove unbound lipoproteins. Cells with VLDL-Dil were subsequently stained with BODIPY to visualize neutral lipid droplets and DAPI to label nuclei. Coverslips were mounted, and fluorescence images were acquired using an epi fluorescence microscope (EVOS M5000) for analysis of VLDL-Dil uptake and lipid droplet distribution. Cells with unlabeled VLDL were harvested for RNA extraction.

### Ex Vivo VLDL Uptake Assay

Immediately following euthanasia, interscapular BAT was rapidly excised and briefly rinsed in ice-cold PBS to remove residual blood and debris. Tissue samples were minced into approximately 1–2 mm explants and transferred to culture wells containing a serum– free culture medium. Explants were allowed to equilibrate for approximately 30 minutes prior to incubation with fluorescently labeled VLDL-Dil (10μg/ml). VLDL uptake was performed as previously described ^35^. Following incubation, explants were washed thoroughly with PBS to remove unbound VLDL-Dil, and fluorescence intensity was measured using a fluorometer to quantify VLDL uptake.

### Adipokine Array

Plasma adipokine profiles were analyzed using the Proteome Profiler™ Mouse Adipokine Array Kit (R&D Systems, Minneapolis, MN, USA; Catalog #ARY013) according to the manufacturer’s instructions. The membrane-based antibody array simultaneously detects the relative expression of 38 adipokines. For both the single binge and acute on chronic treatment experiments, 100μL of plasma from each mouse (n = 8 per group) was used for the assay. Plasma samples were incubated with the supplied biotinylated detection antibody cocktail and then applied to nitrocellulose membranes pre-spotted in duplicate with capture antibodies. Following overnight incubation at 4°C, the membranes were washed and incubated with streptavidin-horseradish peroxidase (HRP). Bound proteins were detected using chemiluminescent detection reagents according to the manufacturer’s protocol, and chemiluminescent signals were captured using a chemiluminescence imaging system. Array images were analyzed by densitometry using ImageJ software. Signal intensities were normalized to the positive control spots on each membrane after background subtraction. Relative adipokine expression levels were compared between treatment groups. Because the assay is semi-quantitative, results are presented as relative changes in protein abundance rather than absolute concentration.

### RT-qPCR

Total RNAs from liver, BAT, and bADs were extracted using RNA Stat60 (Tel-Test) or Trizol (Invitrogen, Cat# 15596026). Complementary DNA was synthesized using the iScript Advanced cDNA Synthesis Kit for RT-qPCR (Bio-Rad). qPCR was performed using a Bio-Rad sequence detection system (Bio-Rad). 18s and GAPDH were used as internal controls for liver, BAT and bADs. The relative amounts of all mRNAs were calculated using the ΔΔCT assay. The mouse primer sequences are shown in Table 2.

**Table 2.**
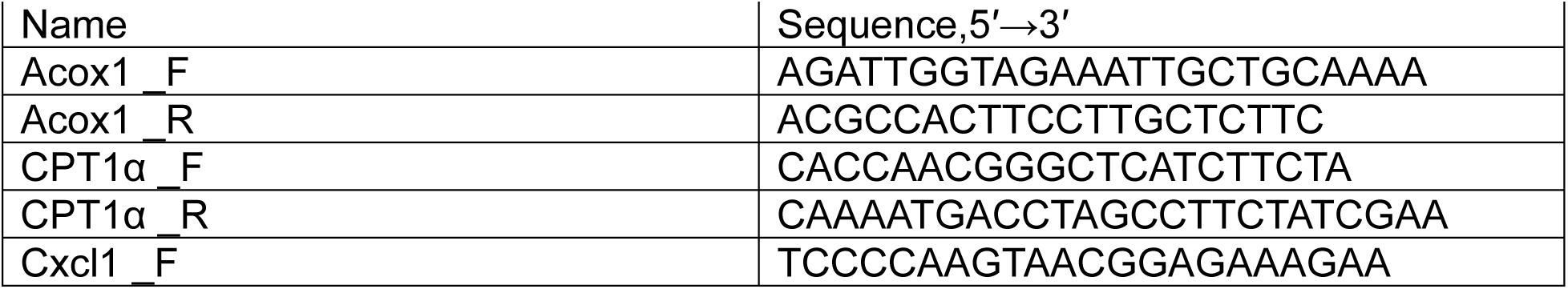
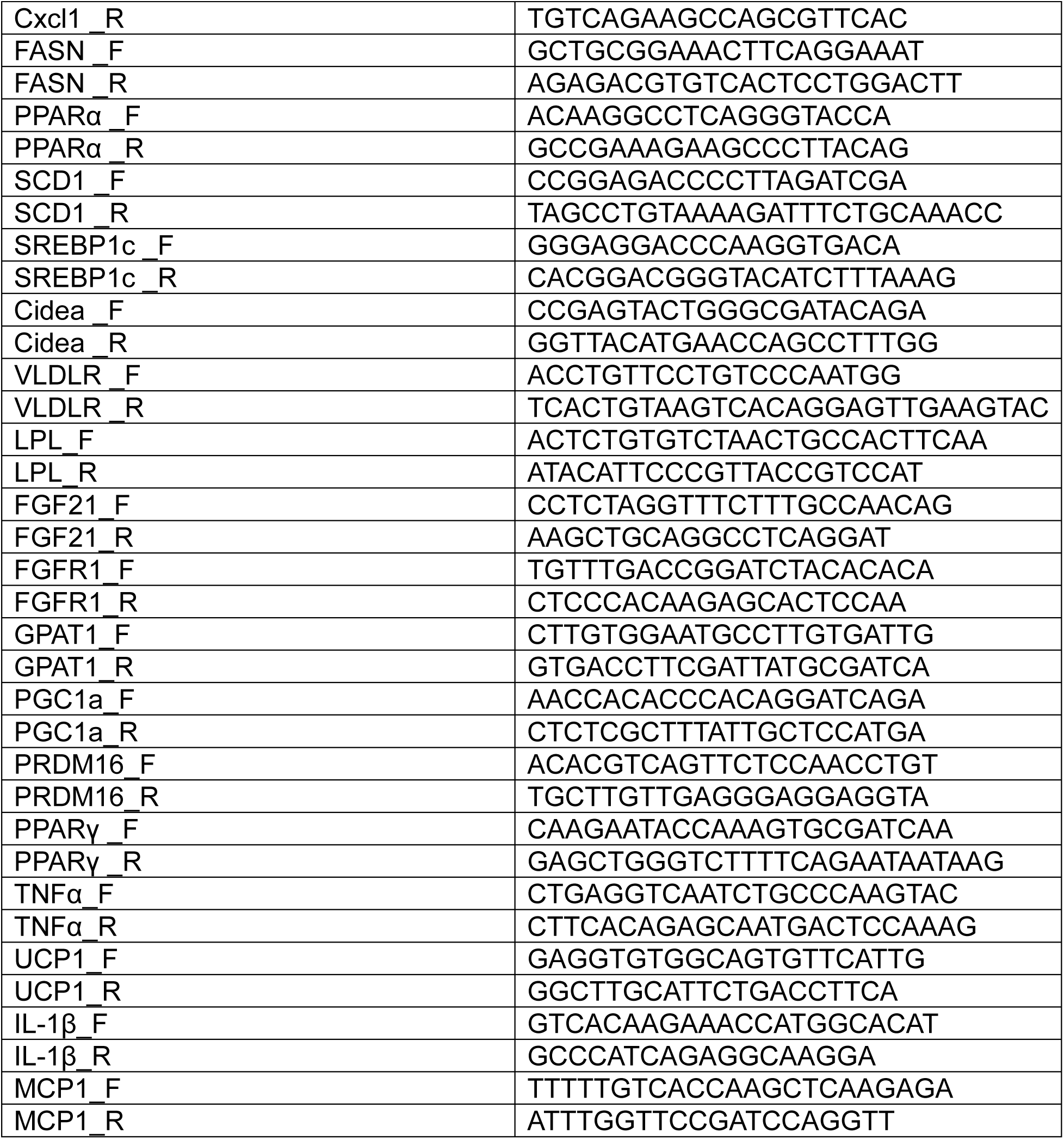
List of Primers.

## Statistical Analysis

Data are expressed as mean ± standard error of the mean (SEM). Statistical analysis was performed using Student’s *t*-test for experiments comparing 2 groups. For studies with 3 or more groups, a 2-way analysis of variance (ANOVA) was used (GraphPad Prism). *P* <.05 is considered significant.

## Acknowledgements

We thank Dr. Kyung Cheul Shin for technical help related to VLDL uptake assay. This work was partially funded by The University of Texas at Dallas Office of Research and Innovation through the New Faculty Research Symposium Grant Program.

---

**Supplementary Figure 1.**
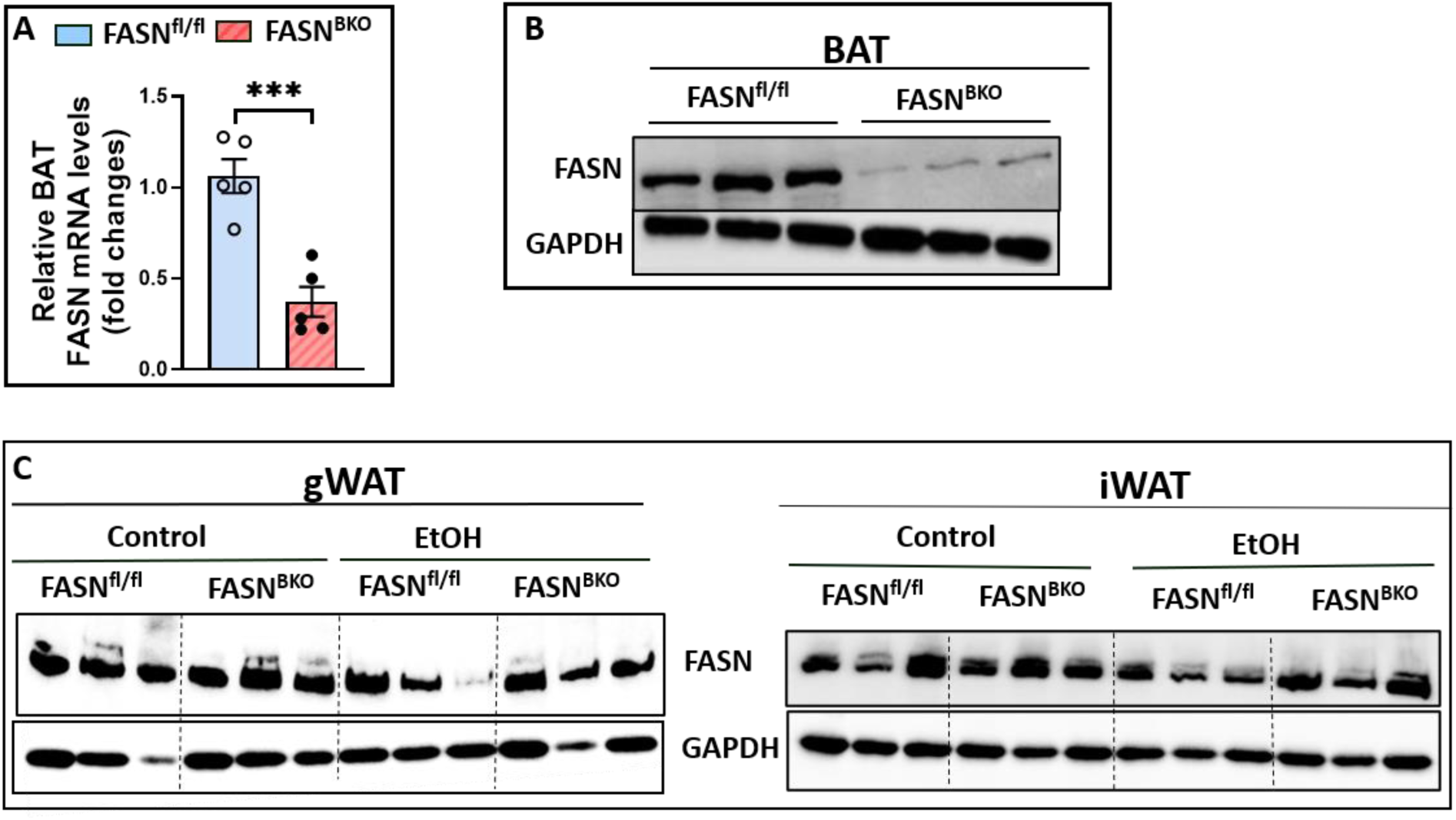
FASN^BKO^ mice exhibit significantly reduced FASN expression in BAT. A) qPCR analysis of FASN mRNA levels in the BAT of chow-fed mice (n=5). (B) Western blotting of FASN and GAPDH in the BAT of chow-fed mice. (C) Male FASN^fl/fl^ and FASN^BKO^ mice were subjected to a single alcohol binge (EtOH, 5 g/kg BW) or isocaloric maltose dextrin (control). Western blotting of FASN and GAPDH in gWAT and iWAT. Data are expressed as the mean ± SEM. ***P <.001.

**Supplementary Figure 2.**
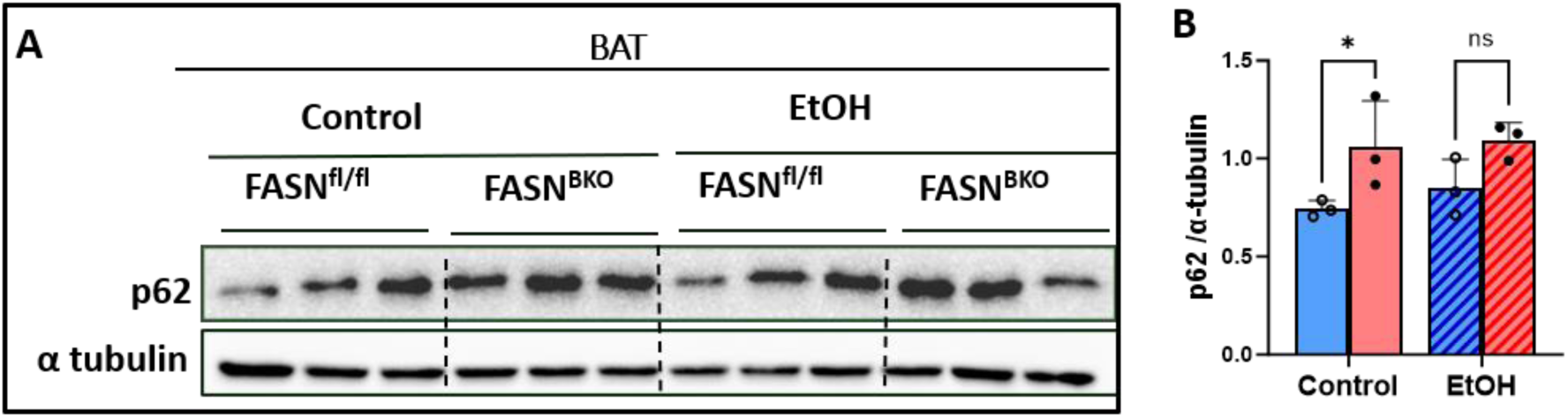
FASN inhibition in BAT increases p62 protein expression. FASN^fl/fl^ and FASN^BKO^ male mice were subjected to a single alcohol binge (EtOH, 5 g/kg BW) or isocaloric maltose dextrin (control) via oral gavage. (A) Western blotting of p62 and α tubulin in BAT. (B) Densitometry quantification of p62 expression from A (n=3). Data are expressed as the mean ± SEM. *P <.05.

**Supplementary Figure 3.**
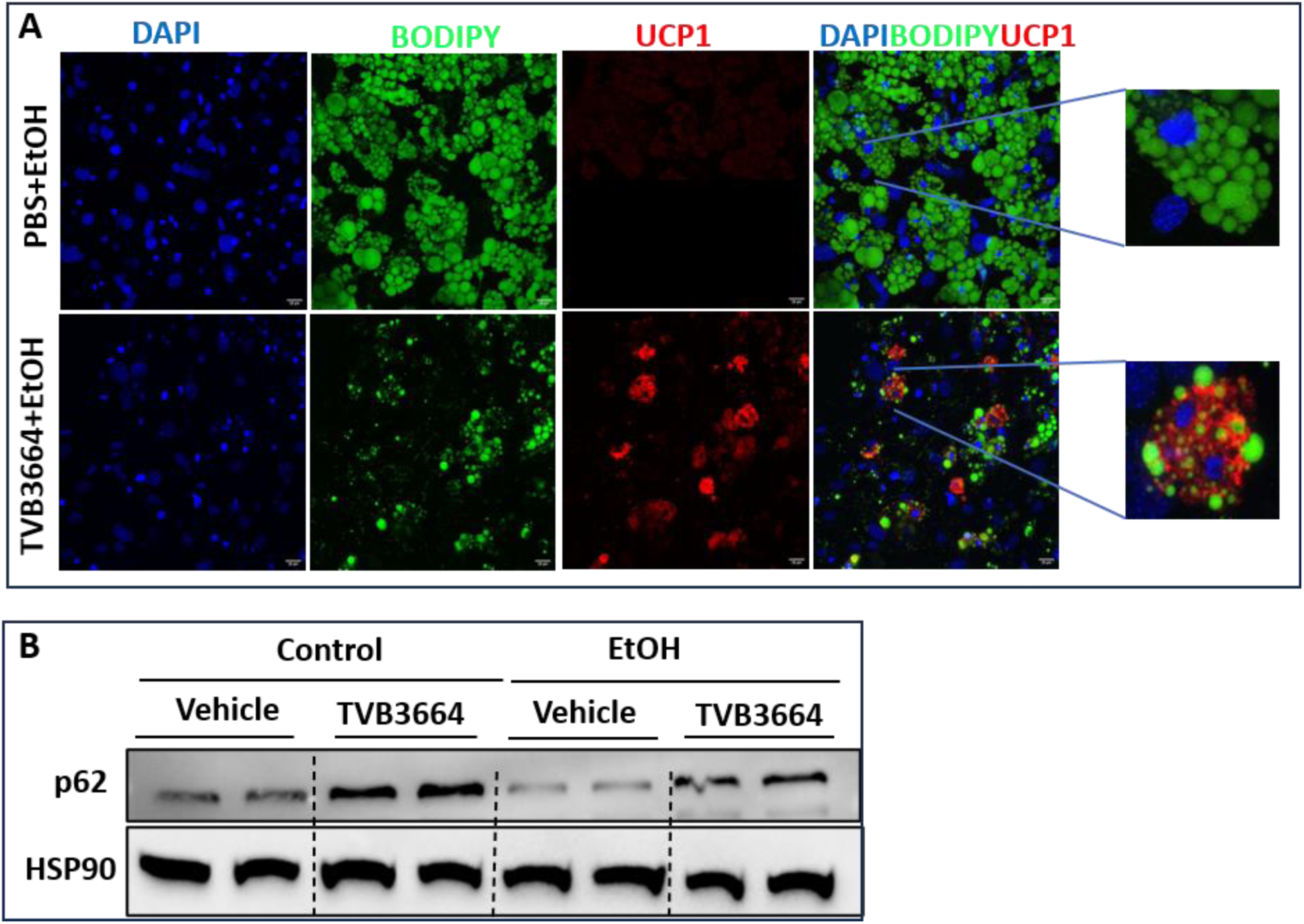
Pharmacological FASN inhibition in bADs increases UCP1 and p62 expression. Differentiated bADS were treated with TVB3664 or vehicle in the presence or absence of 35µM (A) or 100µM EtOH (B). (A) Immunofluorescent staining of UCP1 and BODIPY in bADS. (B) Western blotting of p62 expression in bADs.

**Supplementary Figure 4.**
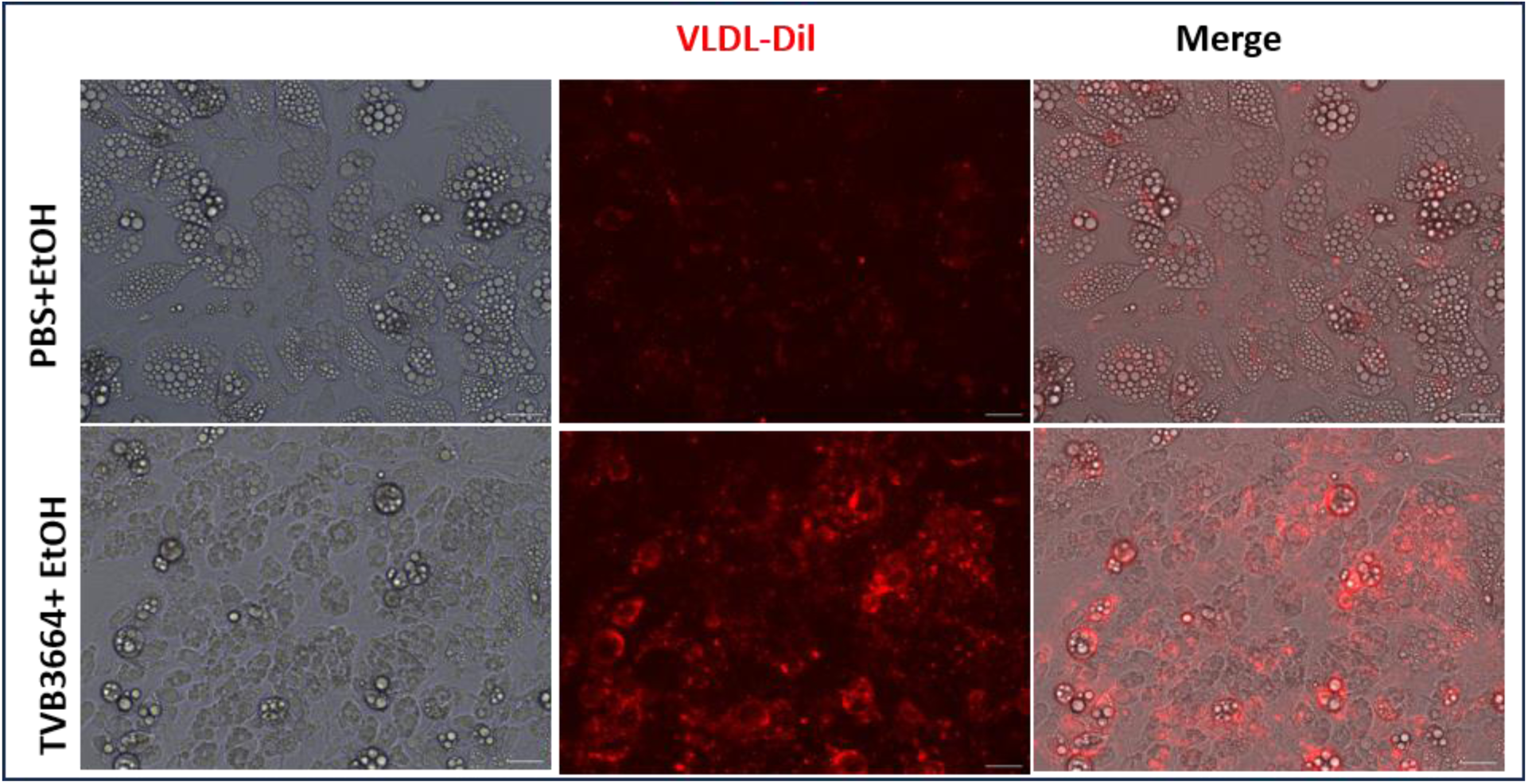
FASN inhibition enhances VLDL uptake in differentiated bADs. bADs were treated with TVB3664 in the presence or absence of 35μM EtOH. Representative images of VLDL-Dil fluorescence in cultured bADs.

**Supplementary Figure 5.**
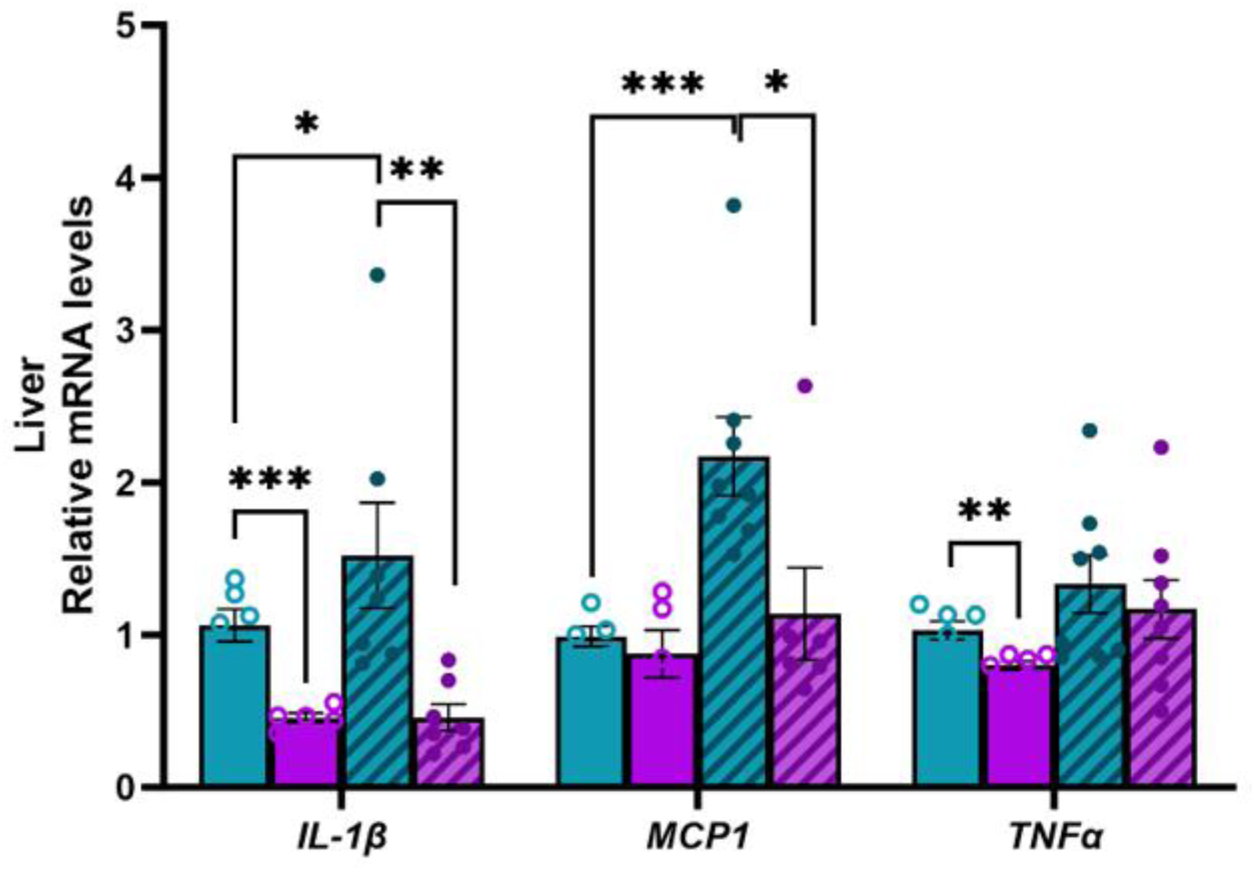
FASN inhibition in BAT rescues acute-on-chronic alcohol-induced hepatic inflammation in mice. qPCR analysis of hepatic expression of IL-1β, MCP1 and TNFα (n=5-8). Data are expressed as the mean ± SEM.*P <.05, **P <.01, ***P <.001.

**Supplementary Figure 6.**
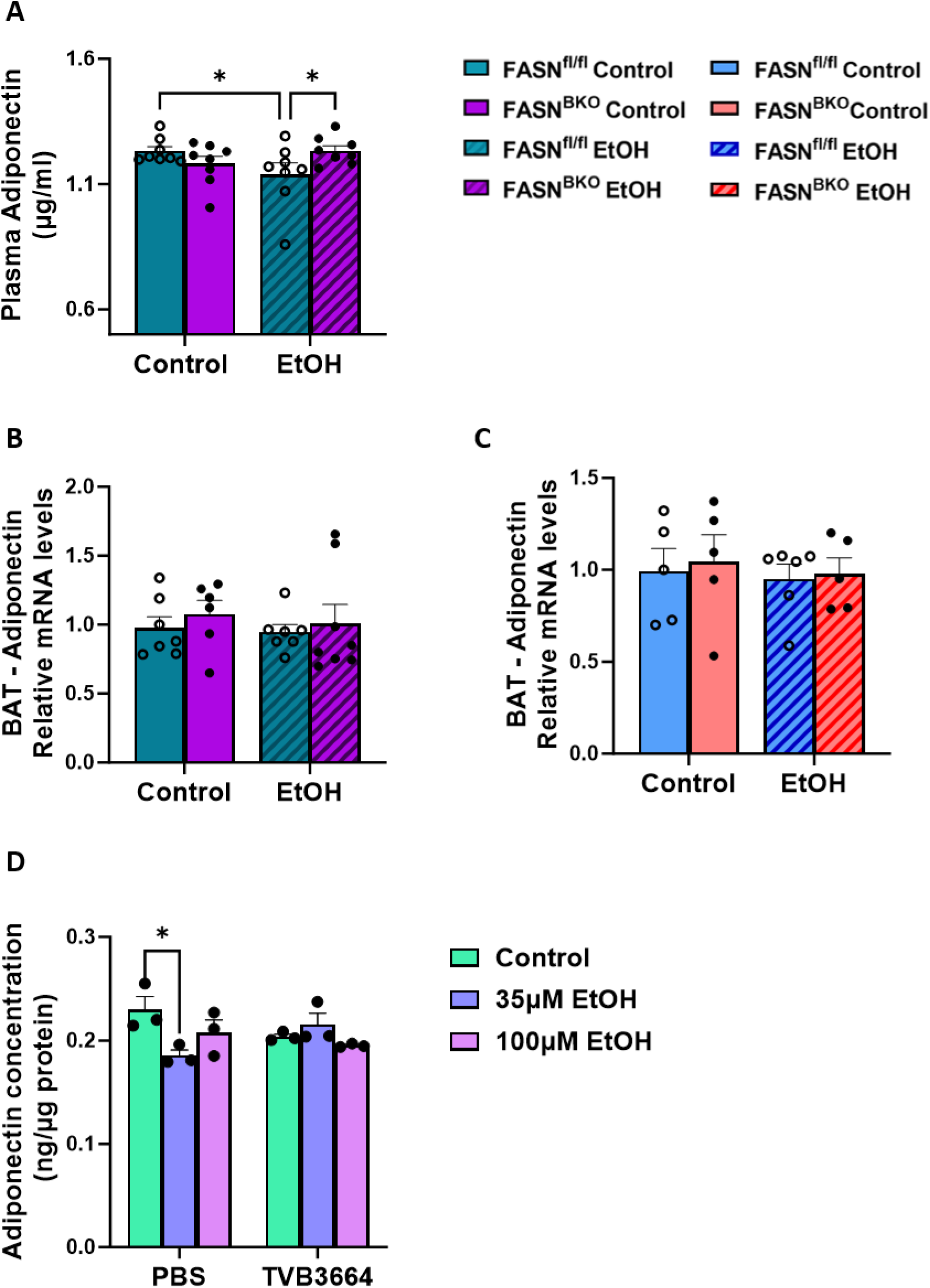
Altered plasma adiponectin levels are observed in acute-on-chronic alcohol-fed FASN^BKO^ mice. (A) Plasma adiponectin levels in mice exposed to acute-on-chronic alcohol feeding model (n= 8). (B-C) qPCR analysis of adiponectin expression in BAT of mice treated with either binge drinking (B) or acute-on-chronic alcohol (C). (D) Adiponectin levels in culture media of differentiated bADs (n= 3). Data are expressed as the mean ± SEM. *P <.05.

